# Adhesion-mediated transition to a mesenchymal-like, fan-shaped migration mode in *Dictyostelium discoideum*

**DOI:** 10.64898/2026.05.03.722454

**Authors:** Gen Honda, Hidenori Hashimura, Satoshi Kuwana, Tomoko Adachi, Daisuke Imoto, Toyoko Sugita, Mitsuru J. Nakamura, Kie Hayashi, Shoko Fujishiro, Mina Fujishiro, Nao Shimada, Satoshi Sawai

**Author notes:** Corresponding author: Satoshi Sawai Department of Basic Science, Graduate School of Arts and Sciences, The University of Tokyo, 3-8-1 Komaba, Meguro-ku, Tokyo 153-8902, Japan. **Author Contributions:** G.H. planned and performed all experiments and image analyses. G.H., H.H., S.K, T.A., T.S., M.J.N, K.H., S.F. and M.F. constructed plasmids and strains. D.I. analyzed cell morphology by machine learning-based classification program. G.H. and S.S. interpreted data and wrote the manuscript. **Competing Interest Statement:** None. **Classification:** Biological Sciences / Cell biology.

## Abstract

Cells migrate with varying degrees of polarization and directional persistence as exemplified by epithelial, mesenchymal and amoeboid cell types. Depending on the physiological and developmental context, these states are often interchangeable, reflecting the plastic and adaptive nature of the cytoskeleton. However, general principles governing such motility-mode transitions remain poorly established, and it is unclear whether they apply to non-metazoan cells. Here, we report previously overlooked features of the amoebozoan *Dictyostelium discoideum,* demonstrating that it undergoes pronounced adhesion-dependent changes in both motility and morphology. Unlike the well-known pseudopodia-rich forms observed on weakly adhesive surfaces, cells on highly adhesive substrates adopt fan-shaped morphologies reminiscent of cultured mesenchymal cells. These cells are characterized by lamellipodia-like protrusions enriched in the SCAR/WAVE complex, large focal adhesion-like plaques, F-actin-independent front-rear gradients of Ras/Rap activity. Furthermore, they exhibit a marked increase in cortical stiffness dependent on F-actin, talins, and the RhoA homolog RacE. Their high directional persistence depends on the persistent localization of the SCAR/WAVE complex, talin-mediated substrate anchoring, and RacE-dependent stabilization of the cell rear. We propose that adhesion-engaged remodeling of cell polarity and cortical mechanics is an evolutionarily ancient feature that predates the specialization of adhesion receptors.

## Introduction

Cell shape changes and migration underpin a wide range of cellular activities, from amoeboid foraging to embryonic development, tissue repair, immune surveillance and cancer metastasis. For crawling cells to move forward, protrusion and retraction of the cell membrane together with cell-substrate adhesion must be tightly coordinated in space and time. At the leading edge, dendritic actin networks are formed by the Arp2/3 complex and its activators, the WASP family proteins, which are widely conserved among eukaryotes, particularly in organisms that migrate via membrane deformation (1, 2). These actin nucleation factors generate a spatially expanding F-actin network that produces propulsive forces through interactions with substrate-anchored adhesion complexes. The shape, persistence and orientation of membrane protrusions are directly linked to migratory behaviors, including cell speed and directional persistence (3). Migrating cells adopt a wide spectrum of shapes that can be broadly classified into several types. Immune cells and free-living amoebae are often categorized as fast-migrating cells; they rapidly extend pseudopods, exhibit weak substrate adhesion, and form small, transient adhesive clusters known as ventral adhesion foci (4). In contrast, mesenchymal and epithelial cells migrate more slowly and are characterized by thin, broad lamellipodia supported by the Arp2/3 complex and large, stable focal adhesion complexes, representing the other end of the spectrum of adhesion-dependent migratory behaviors. In these cells, large focal adhesions form behind lamellipodia through Rap1-mediated activation of talin (5), in concert with F-actin-dependent mechanical activation of the vinculin-talin complex.

Cell shape and motility can change drastically in response to the physicochemical environment. Such phenotypic changes can be rapidly induced by experimental perturbations, including pharmacological inhibition, optogenetic activation, modulation of substrate adhesion, and physical confinement (6–11). For instance, certain cancer cells adopt distinct morphologies depending on culture conditions, exhibiting a mesenchymal phenotype in adherent culture and a blebby phenotype in suspension (6). Under conditions of low adhesion and physical confinement, cultured cancer cells can switch to a fast-migrating mode (7). Transitions between distinct migration modes are thought to confer energetic advantages (12, 13) and drug resistance during cancer invasion in 3D matrices (14–16). These transitions, often referred to as the mesenchymal-to-amoeboid transition in the context of cancer metastasis, are thought to be governed by the balance between Rac and Rho signaling, which in turn influences the balance between Arp2/3-mediated actin assembly and actomyosin contractility (6, 10, 17, 18). The amoeboid motility has also been characterized in fibrosarcoma cells by the loss of integrin clusters (16) and transcriptional changes such as increased expression of RhoA and the PI3K/PKB-related pathway (19).

With the exception of pathogenic *Entamoeba* species which have been shown to exhibit fibronectin-induced motility transitions (20), how cell-substrate adhesion governs migration and polarity in non-metazoan cells remains largely unexplored. In the genetically tractable *Dictyostelium*, previous studies have suggested that the shape of cell protrusions is strongly dependent on substrate adhesion (21–24). While *Dictyostelium* lacks integrins, it possess five integrin β-like Sib proteins (25, 26), all of which interact with the talin homolog TalA (25, 27). As free-living amoebae, *Dictyostelium* cells can adhere to various substrates, including borosilicate, polystyrene, mica, and SU-8 polymer (4, 28), enabling migration and phagocytosis without requiring specific surface ligands (25, 29). The traction forces generated by these cells are on the order of 0.1–1 nN (30, 31), comparable to estimated van der Waals forces at the cell-substrate interface (32). Substrate hydrophobicity and surface charge strongly influence adhesion strength (4), with adhesion receptors and the glycocalyx cooperatively contributing to EDL-DLVO (Electrical Double Layer-Derjaguin-Landau-Verwey-Overbeek)-type forces (33). Under the microscope, TalA is observed in adhesion foci (34), and similar focal accumulations have been observed for the other talin homolog TalB (35) as well as vinculin homologues VinA and VinB (35, 36), paxillin homolog PaxB (37), myosin VII (38), α-actinin (39). Rap1 serves as a key regulator of *Dictyostelium* cell-substrate adhesion (40) by allosterically activating TalB (41) and interacting with additional adhesion-related proteins, such as Ser/Thr-kinase Phg2 (21, 42), Ste20-familiy kinase KrsB (43), and the IQGAP-related protein IqgC (44). While focal adhesion kinase (FAK) and Src family kinases are exclusive to metazoans (45), these studies indicate that *Dictyostelium* adhesion relies on small, short-lived adhesion foci that incorporate key ancestral components of the metazoan focal adhesion machineries.

Here, we employed substrate modifications to facilitate cell-substrate attachment and systematically analyzed polarization and migration of *Dictyostelium discoideum* cells on uniform substrates with different adhesion strengths. We show that increased adhesion drastically alters cell shape and migration trajectories, yielding mesenchymal-like morphologies. Enhanced substrate adhesion promotes leading-edge enrichment of the SCAR complex and focal adhesion-like plaques, stabilizes the front-rear gradient of Rap1-GTP, and increases cellular stiffness. This spreading-associated polarization and stiffening require talin and the RhoA homolog RacE, suggesting that adhesion-triggered activation of contractility and cell polarization represents an evolutionarily ancient feature that predates the specialization of adhesion receptors.

## Results

### Enhanced substrate attachment induces fan-shaped migration of *Dictyostelium* cells

To facilitate cell-substrate attachment, borosilicate glass coverslips were coated with wheat germ agglutinin (WGA) lectin. WGA selectively binds N-acetylglucosamine residues on the cell surface. Under this condition, the majority of aggregation-stage AX4 cells attached to WGA-coated substrates migrated with a flattened, fan-shaped or laterally elongated shape, reminiscent of fish keratocytes (Fig. 1A, “WGA”), in stark contrast to the typically observed linear shape on uncoated coverslips (Fig. 1A, “Uncoated”). To investigate whether this fan-shaped migration was specific to WGA, we tested two additional surface coatings: poly-L-lysine (PLL) and Cell-tak (CTK). PLL is a cationic polymer that promotes adhesion between negatively charged membranes and glass substrates, whereas CTK is a polyphenolic protein mixture extracted from *Mytilus edulis* that acts as a nonspecific adhesive to a wide range of materials (46). Fan-shaped migration was also observed on both PLL- and CTK-coated substrates (Fig. 1A, “PLL” and “CTK”), although the occurrence was lower (Fig. 1B). At concentrations above 10 µg/mL PLL and 100 µg/mL CTK, cells frequently ruptured immediately after spreading, which may result from impaired shedding of adhesion receptors at the trailing edge (24). Nevertheless, these observations indicate that the observed changes in cell shape are independent of the specific ligand.

**Figure 1.**
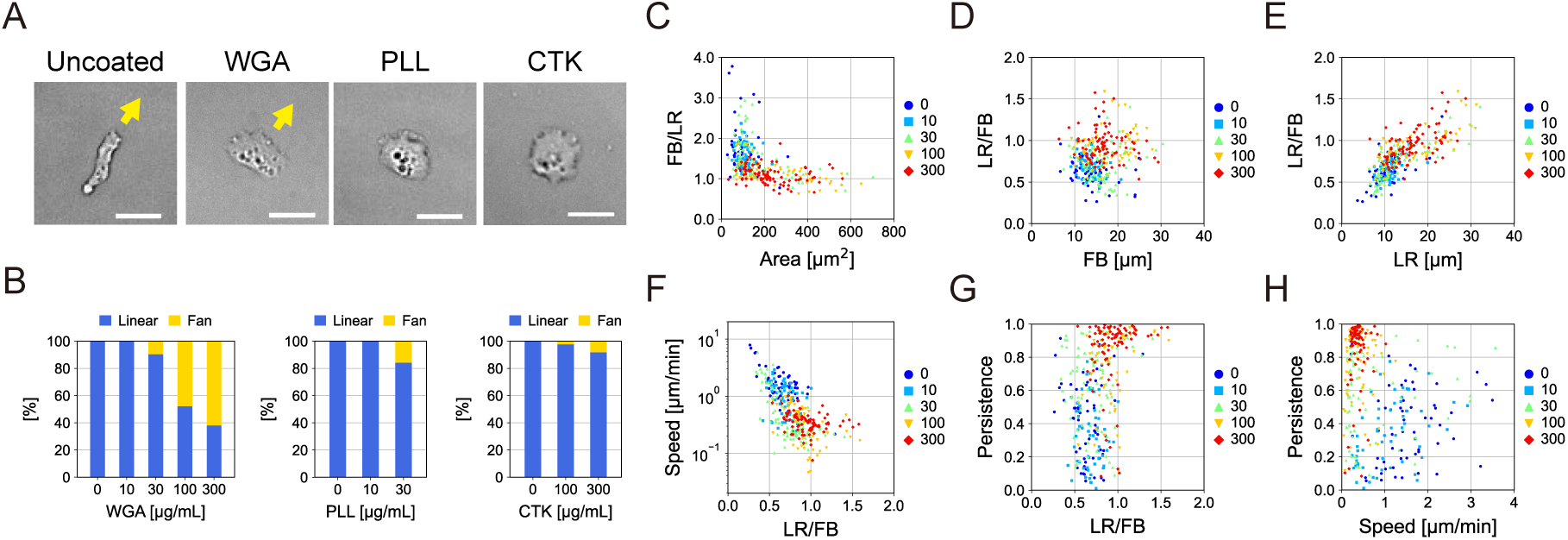
Enhanced cell-substrate attachment induces fan-shaped migration of *Dictyostelium* cells. (**A**) Representative snapshots of aggregation-stage AX4 cells on uncoated and WGA-, PLL-, CTK-coated borosilicate coverslip. Yellow arrows indicate direction of migration. Scale bars, 20 µm. (**B**) Percentage of linear- and fan-shaped cells on WGA-, PLL-, CTK-coated substrates. N > 32 cells each. (**C**–**H**) Scatter plots of cell shape and motility on WGA-coated substrates. Legend, WGA concentration (µg/mL). N = 54, 74, 78, 62, 83 cells. Each dot represents individual cells. FB: front-to-back length of cells. LR: left-to-right width of cells. (**C**) Cell area versus aspect ratio FB/LR. (**D**–**E**) FB or LR versus inverse aspect ratio LR/FB. (**F**–**G**) LR/FB versus cell speed and persistence. (**H**) Cell speed versus persistence.

To quantitatively characterize cell shape and migratory behavior, we performed time-lapse imaging of GFP-Lifeact-expressing cells migrating on WGA-coated substrates. Two shape features were extracted for each cell trajectory (30–50 min): the spreading area and the ratio of front-back length to left-right width (FB/LR), averaged over time. At low WGA density (10 µg/mL), cell shape was indistinguishable from that on uncoated substrates, exhibiting elongated anterior-posterior shapes, with FB/LR values between 1.0 and 4.0 and spreading areas below 200 µm^2^ (Fig. 1C, “0” and “10”). In contrast, at high WGA densities (100–300 µg/mL), FB/LR values ranged from 0.5 to 2.0 and spreading areas ranged from 100 to 800 µm^2^ (Fig. 1 C, “100” and “300”), corresponding to circular (FB/LR ∼ 1) or laterally elongated (FB/LR < 1) shapes. At an intermediate WGA density (30 µg/mL), cells occupied both shape regimes (Fig. 1 C, “30”). Across the range of WGA concentrations (0–300 µg/mL), the average front-back length increased by 23.8%, whereas the left-right width increased by 50.9%. Accordingly, the inverse aspect ratio (LR/FB) correlated strongly with changes in lateral width (correlation coefficient = 0.70) but not with front-back length (correlation coefficient = 0.06) (Fig. 1D–E). These results indicate that enhanced substrate attachment preferentially promotes lateral spreading, leading to a breakdown of the typical anterior-posterior aspect ratio. This was also confirmed using a deep learning-based cell shape classifier (3), which projects diverse cell morphologies onto a two-dimensional principal component analysis (PCA) space (Fig. S1A). *Dictyostelium* cells on uncoated substrates localized to the *Dictyostelium*-specific region (Fig. S1B, *left*), whereas those on WGA-coated substrates shifted to regions characteristic of fan-shape morphology (Fig. S1B, *right*).

Cell motility parameters were extracted from centroid trajectories by fitting a persistent random walk model to the mean squared displacement (MSD) (Fig. S1C–D). With increasing WGA density, the cell speed decreased from 3.3 to 0.43 µm/min, whereas the persistence time increased from 3 to 30 min (Fig. S1E). To further relate cell shape to individual migration trajectories, directional persistence was also quantified as the ratio of net centroid displacement to total path length. Cell speed decreased with increasing WGA density and correlated with inverse cell aspect ratio (Fig. 1F). In contrast, persistence increased with WGA density, approaching 1 (Fig. 1G), and was higher in laterally elongated cells (LR/FB > 1) than in cells with other shapes (LR/FB < 1) (Fig. 1G). Consequently, under conditions of strong substrate attachment, cells migrated slowly but with highly directed trajectories (Fig. 1H), resembling migration modes typically associated with integrin-mediated adhesion systems (7).

### Enhanced cell-substrate attachment induces membrane recruitment of the SCAR complex and Arp2/3 dependent lamellipodia-like protrusions

On uncoated substrates, migrating *Dictyostelium* cells typically extended narrow, finger-like pseudopodia (Fig. 2A, Uncoated, *top*). In contrast, on WGA-, PLL-, or CTK-coated substrates, fan-shaped cells formed thin, broad membrane protrusions at the leading edge (Fig. 2A, WGA, PLL and CTK, *top*, yellow arrows), reminiscent of lamellipodia in metazoan cells. Three-dimensional reconstruction of fan-shaped cells showed that F-actin, visualized with GFP-Lifeact, was enriched along the edges of the cell-substrate interface (Fig. S2A). When cells were sandwiched between WGA-coated substrates, sheet-like protrusions formed on both sides of the cell (Fig. S2B), but not on uncoated substrates (Fig. S2C). These indicate that sheet-like protrusions are generated at cell-substrate interfaces. Total internal reflection fluorescence (TIRF) microscopy revealed that the SCAR complex subunit HSPC300 fused to GFP (Fig. 2A, WGA, PLL and CTK, *bottom*) and GFP-Arp2 (Fig. S2D, WGA) were enriched at the distal edge of sheet-like protrusions. F-actin also displayed a laterally expanded distribution along the leading edge (Fig. 2A, WGA, PLL and CTK, *middle*). These localization patterns differed markedly from those observed at the tips of finger-like pseudopodia on uncoated substrates (Fig. 2A, Uncoated, *middle* and *bottom*). Pharmacological inhibition of Arp2/3 with CK666 abolished the sheet-like protrusions, induced cell rounding, and completely arrested migration (Fig. S2E–F). Furthermore, fan-shaped, highly persistent migration was lost in *pirA*^−^ cells, which lack the SCAR complex subunit PIR121, and was rescued by re-introduction of PIR121 (Fig. S2G–H). Together, these results indicate that fan-shaped migration on adhesive substrates depends on the SCAR complex and Arp2/3-mediated actin nucleation.

**Figure 2.**
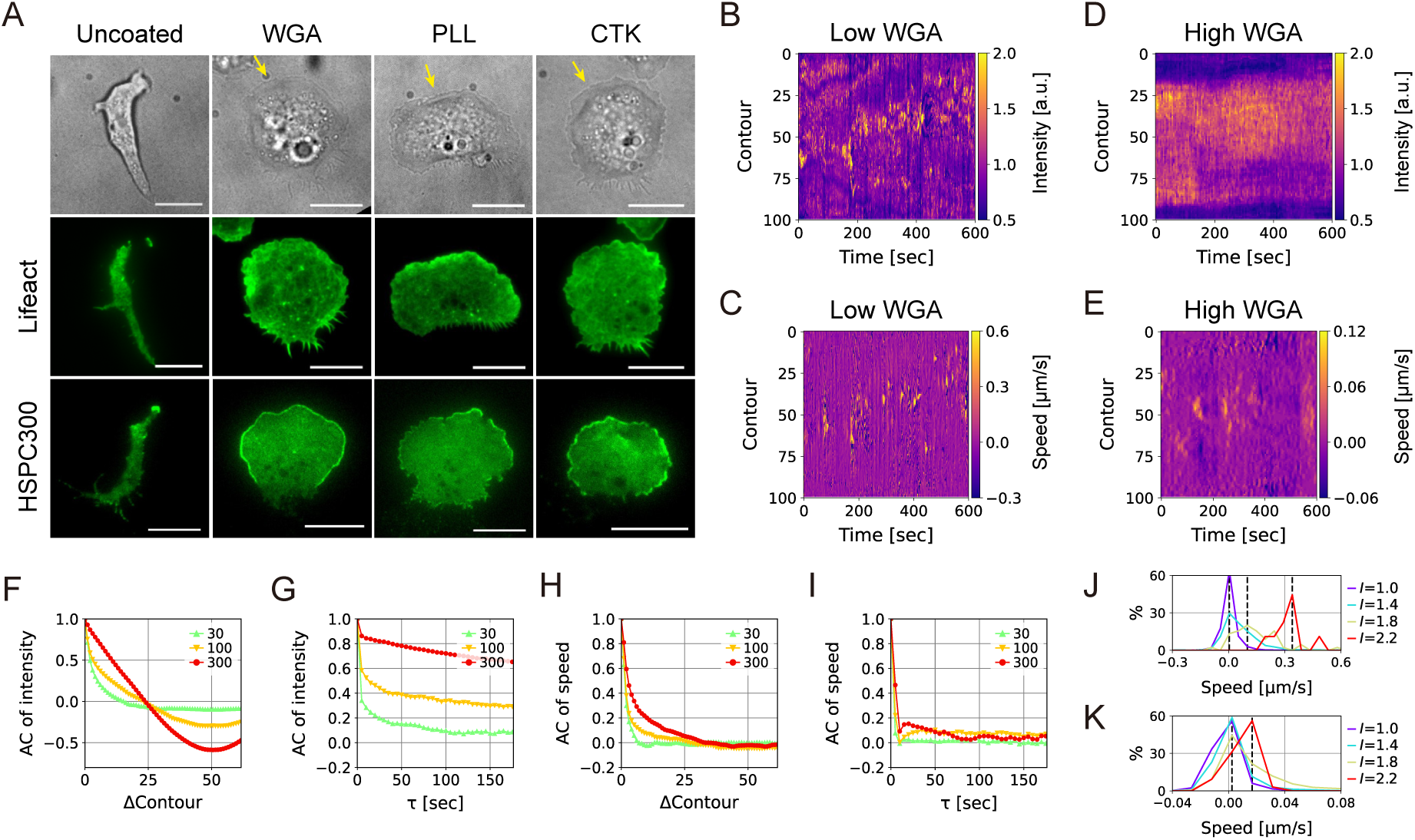
Strong substrate adhesion induces persistent localization of the SCAR complex at the edge of lamellipodia-like protrusions. (**A**) Transmitted-light (*top*) and TIRF images of GFP-Lifeact (*middle*) and HSPC300-GFP (*bottom*) in cells attached to uncoated and WGA-, PLL- and CTK-coated substrates. Scale bars, 10 µm. (**B**–**E**) Spatiotemporal profiles of the SCAR complex and membrane protrusions. (**B**, **D**) Fluorescence intensity of HSPC300-GFP near the edge, normalized to the value at the proximal region. (**C**, **E**) Protrusion speed. Substrates were coated with 30 µg/mL WGA in **B**–**C**, and 300 µg/mL in **D**–**E**. (**F**–**I**) Autocorrelations of HSPC300-GFP intensity in space (**F**) and time (**G**). Autocorrelations of protrusion speed in space (**H**) and time (**I**). Legend, WGA concentration (µg/mL). N = 4 cells each. (**J**–**K**) Histograms of local protrusion speed formed at the region of normalized HSPC300-GFP intensity *I* = 1.0, 1.4, 1.8 and 2.2. Substrates were coated with 30 (**J**) and 300 µg/mL WGA (**K**). Broken lines, modal values.

To quantify protrusion dynamics, we applied a boundary tracking method (47) to analyze both membrane displacement and HSPC300-GFP intensity along the cell contour. At low WGA density, HSPC300-GFP appeared as narrow (∼ 10% of the total contour), sporadic, and transient bursts (Fig. 2B), coinciding with localized membrane protrusions (Fig. 2C). In contrast, at high WGA density, HSPC300-GFP intensity was broadly distributed along the boundary (>50% of the contour) and persisted over time (Fig. 2D). Despite this broad and stable HSPC300 localization, membrane protrusions remained relatively narrow and short-lived (Fig. 2E). Consistent with these observations, both spatial and temporal autocorrelations of HSPC300-GFP intensity increased markedly at higher WGA density (Fig. 2F–G). In contrast, the corresponding autocorrelations of membrane protrusions increased only modestly (Fig. 2H–I). To enable quantitative comparison across conditions, autocorrelation curves were fitted with exponential functions to extract characteristic spatial and temporal scales. At 30 µg/mL WGA, protrusion width was characterized by a single spatial scale corresponding to 1.8% of the cell contour (Fig. S3A–B). At 300 µg/mL WGA, two spatial components were required, corresponding to widths of 2.0% and 10.7% of the contour (Fig. S3A–B), indicating that the coexistence of protrusions with distinct lateral extents, as also apparent in the spatiotemporal heatmaps (Fig. S3C). In contrast, temporal autocorrelations were well described by a single timescale at all WGA densities, with protrusion lifetimes increasing only modestly from 1.7 to 4.7 s (Fig. S3D–E). These results indicate that fan-shaped cells migrate via a stepping mode of protrusion (30, 48), rather than smooth, continuous gliding as observed in fish keratocytes.

To further investigate the relationship between the SCAR complex localization and membrane protrusion, we analyzed 2D histograms relating local protrusion speed to HSPC300-GFP intensity, normalized to the proximal cell region. At low WGA density (Fig. 2J; Fig. S2F), protrusion speed was symmetrically distributed around zero at normalized intensity *I* = 1.0, and became increasingly skewed toward positive values as intensity increased. The modal protrusion speed exceeded zero at *I* = 1.8 (0.10 µm/s) and further increased to 0.34 µm/s at *I* = 2.2. In contrast, at high WGA density (Fig. 2K; Fig. S2G), protrusion speed remained symmetrically distributed up to *I* = 1.4, became skewed at *I* = 1.8, and the modal protrusion speed increased above zero at *I* = 2.2 (0.017 µm/s). These analyses indicate that enhanced cell-substrate attachment markedly increases the extent and persistence of SCAR complex accumulation at the leading-edge, while membrane protrusion requires higher local levels of SCAR activity under strongly adhesive conditions.

### Enhanced substrate adhesion does not recruit WASP

In addition to the SCAR complex, WASP is a major activator of the Arp2/3 complex. Because SCAR and WASP can partially compensate for each other during cell migration (49, 50), we asked which of these nucleation promoting factors is recruited. On WGA-coated substrates, HSPC300-GFP was localized broadly along the front edge in *wasA*^−^, which lacks WASP-A, the only *Dictyostelium* WASP homolog that localizes to the leading edge (2), as well as in its parental strain AX2 (Fig. S4A–B), indicating that recruitment of the SCAR complex to the edge occurs independently of WASP. Conversely, in *pirA*^−^ cells, WASP-A localized to narrow regions of the edge (Fig. S4D), in contrast to broad edge localization in wild-type cells (Fig. S4C). This broad and persistent edge localization of WASP-A was restored in *pirA*^−^ by re-introduction of PIR121 (Fig. S4E). We note that although SCAR itself is largely absent in *pirA*^−^, another SCAR complex subunit, Nap1, remains expressed in this mutant (51). Similar to HSPC300, Nap1 was enriched at the edge in wild-type cells on WGA-coated substrates (Fig. S4F), whereas it was absent in *pirA*^−^ (Fig. S4G). Edge enrichment of Nap1 in *pirA*^−^ cells was rescued by PIR121 expression (Fig. S4H), confirming that recruitment of the SCAR complex to the cell edge requires proper subunit assembly. Together, these results indicate that enhanced cell-substrate attachment selectively facilitates edge localization of the SCAR complex, but not WASP.

Similar relationships were observed when cortical F-actin was disrupted. Treatment with latrunculin A (LatA) on weakly adhesive substrates induced the formation of numerous membrane-associated puncta of both SCAR complex subunits and WASP-A (Fig. S5A), as previously reported for HSPC300 (22, 52). LatA treatment of *wasA*^−^ and *pirA*^−^ cells recapitulated the patterns observed in untreated cells; HSPC300 formed membrane-associated puncta in *wasA*^−^ (Fig. S5B), whereas in *pirA*^−^ cells, only WASP-A, but not SCAR complex subunits, was targeted to the membrane (Fig. S5D). Notably, when wild-type and *wasA*^−^ cells were treated with LatA on WGA-coated substrates, SCAR complex subunits were predominantly enriched at the cell edge and along membrane nanotubes extending around the cells (Fig. S5A, “WGA” and S5C). In contrast, WASP-A was not enriched at these sites; instead, its puncta were uniformly distributed across the membrane in both wild-type (Fig. S5A) and *pirA*^−^ cells (Fig. S5E). These membrane nanotubes formed upon disruption of cortical actin (53), likely driven by substrate adhesion-mediated pulling forces and possibly aided by BAR domain-mediated membrane deformation. In support of this notion, fluorescently tagged IBARa, an I-BAR and SH3 domain-containing protein (54, 55), appeared as membrane-associated puncta in LatA-treated cells on uncoated substrates (Fig. S5F, “Uncoated”), and became enriched along the cell edge and at the tips of membrane nanotubes on WGA-coated substrates (Fig. S5F, “WGA” and S5G). These observations indicate that the edge-specific recruitment of the SCAR complex under strongly adhesive conditions is independent of cortical F-actin integrity.

### Adhesion-induced fan-shaped migration is independent of PI(3,4,5)P3 signaling

The SCAR complex is known to be recruited to the protruding rims of macropinocytic cups, expanding with patches of Ras-GTP and phosphatidylinositol (3,4,5)-trisphosphate (PIP3) (56). To test whether a similar signaling architecture underlies fan-shaped migration on adhesive substrates, we examined the distribution of phosphoinositide markers. On WGA-coated substrates, the PIP3 marker PH_CRAC_-GFP and PH_Akt_-GFP formed discrete, pinosome-like patches at the ventral membrane but were not enriched at the leading edge (Fig. S6A). In contrast, markers for PI(4,5)P2 exhibited distinct distributions: PTEN showed a gradual posterior enrichment, cortexillin I accumulated at the trailing edge and PH_PLCδ_ was uniformly distributed along the ventral surface (Fig. S6A). The phosphatidylserine probe LactC2 likewise did not accumulate at the leading edge. Pharmacological inhibition of PI3K with LY294002, which disrupts macropinocytic patches (57, 58), markedly suppressed cell deformation and migration on uncoated substrates (Fig. S6B, “Uncoated”). In contrast, the same treatment had no significant effect on cell shape or migratory behavior on WGA-coated substrates (Fig. S6B, “WGA” and S6C–D). Consistent with this result, *pi3k1-5*^−^ and *pten*^−^ cells on WGA-coated substrates adopted polarized fan-shape and displayed broad leading-edge localization of HSPC300-GFP (Fig. S6E). These results indicate that fan-shaped migration induced by strong cell-substrate attachment is independent of PIP3 signalling. This adhesion-driven mode of migration is therefore mechanistically distinct from the previously reported fan-shaped cell migration driven by propagating PIP3 patches (57, 59, 60).

### Front-rear gradients of Ras and Rap1 activity are maintained independently of F-actin

The small GTPase Rac1 directly activates the SCAR complex, whereas Ras and Rap1 function as their upstream regulators. To examine the spatial organization of these signaling molecules during adhesion-induced migration, we visualized their active, GTP-bound forms using TIRF microscopy with GFP-RBD_Raf1_ (Ras-GTP), GFP-RBD_RalGDS_ (Rap1-GTP) and GFP-CRIB_PakB_ (Rac1-GTP). On uncoated substrates, all three probes were enriched at the leading edge of migrating cells (Fig. 3A, *left*, “Uncoated”), consistent with canonical front-localized signaling during amoeboid migration. In contrast, on WGA-coated substrates, these probes were broadly distributed across the ventral membrane with a gradual decrease in signal intensity toward the posterior end (Fig. 3A, *center*, “WGA”). Previous studies showed that disruption of F-actin with LatA induces propagating patches of Ras-GTP, Rap1-GTP and PIP3 across the plasma membrane (52, 58, 61), whereas Rac1-GTP becomes uniformly distributed (62). Unexpectedly, when cells on WGA-coated substrates were treated with LatA, front-rear gradients of GFP-RBD_Raf1_ and GFP-RBD_RalGDS_ persisted for more than 30 min (Fig. 3A, *right*, “WGA; +LatA” and 3B, “RBD_Raf1_” and “RBD_RalGDS_”). In contrast, GFP-CRIB_PakB_ exhibited the uniform distribution, occasionally forming bright patches in the central and posterior regions (Fig. 3A, *right*, “WGA; +LatA” and 3B, “CRIB_PakB_”). Time-averaged fluorescence intensity profiles along the cell midline revealed a front-to-rear gradient for RBD_Raf1_, RBD_RalGDS_, and CRIB_PakB_ in untreated cells, whereas PH_Akt_ was depleted at the leading edge and relatively uniform across the central region (Fig. 3C, “Intact”). Following LatA treatment, spatial gradients of RBD_Raf1_ and RBD_RalGDS_ were preserved, with the RBD_RalGDS_ gradient becoming even steeper, whereas the gradient of CRIB_PakB_ was lost (Fig. 3C, “LatA”). These observations indicate that strong cell-substrate adhesion promotes and maintains a front-rear gradient of Ras and Rap1 activity independently of F-actin.

**Figure 3.**
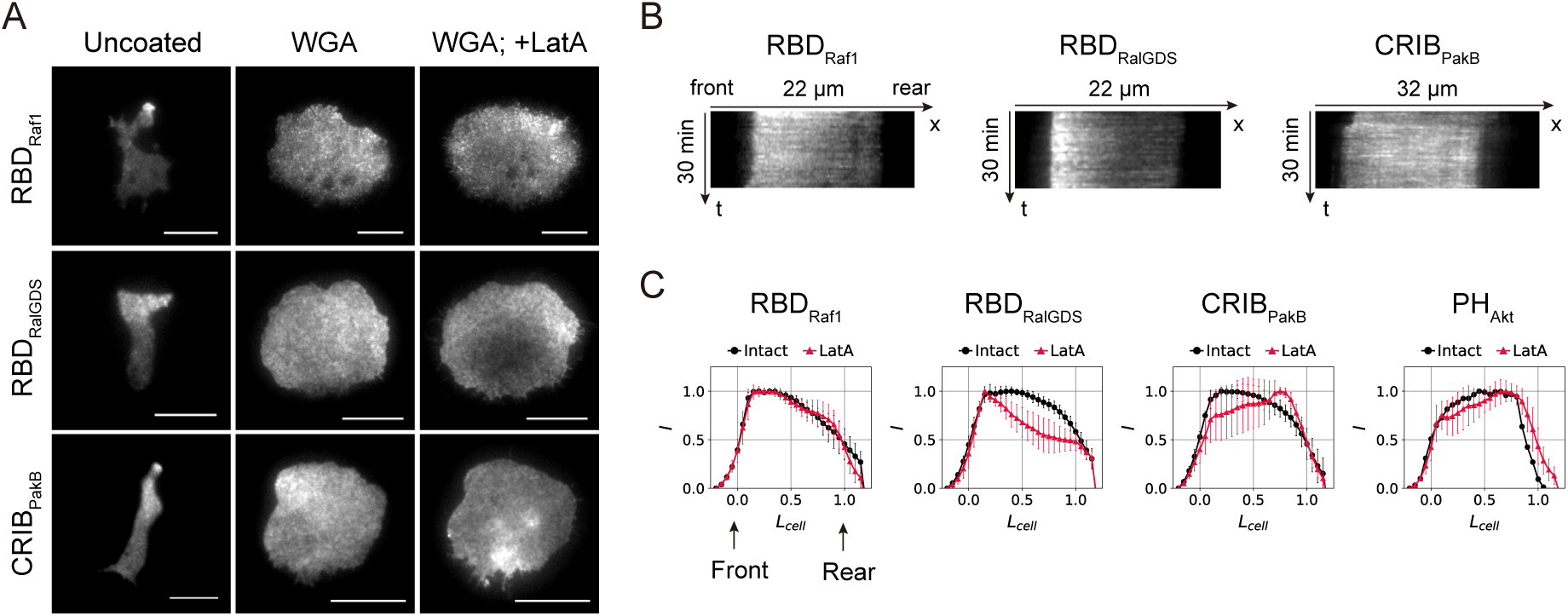
Front-rear gradients of Ras-GTP and Rap1-GTP persist after F-actin disruption. (**A**) TIRF images of GFP-RBD_Raf1_, RBD_RalGDS_-GFP, and CRIB_PakB_-GFP in cells attached to uncoated (*left*) and WGA-coated substrates (*center* and *right*). (*right*) 3 µM LatA-treated. (**B**) Kymographs taken along the midline of cells treated with LatA on WGA-coated substrates. *t* = 5 to 35 min after LatA treatment. (**C**) Front-rear profiles of fluorescent probes (mean ± s.d, N = 4–14 cells). Cell length is normalized so that the leading edge is aligned to *x* = 0 and the trailing edge to *x* = 1.

### Enhanced attachment promotes the formation of focal adhesion-like plaques

Next we analyzed the localization of focal adhesion-related proteins. On WGA-coated substrates, GFP-PaxillinB (PaxB) formed large, long-lived foci near the cell edge (Fig. 4A). These focal adhesion-like plaques required F-actin as they disappeared upon LatA treatment (Fig. 4A, “+LatA”). At low WGA density, GFP-PaxB formed small, transient foci at protruding tips (Fig. S7A), whereas high WGA density induced a dense line of PaxB assemblies along the cell edge, which dynamically remodeled during protrusion-retraction cycles (Fig. S7B–C). These periodic assembly and disassembly dynamics of PaxB paralleled the discontinuous protrusive behavior of fan-shaped migration (Fig. 2E). While undetectable at low WGA density, GFP-VinculinA (VinA) also formed large PaxB-like foci at high WGA density (Fig. 4B). Talin isoforms exhibited distinct localization patterns: TalinA-GFP was distributed as numerous puncta across the ventral surface and showed enrichment toward the trailing edge at high WGA density (Fig. 4C), whereas Neon-TalinB formed fewer ventral foci and preferentially localized to the leading edge (Fig. 4D). Following F-actin disruption, TalinA-GFP partially retained ventral association, whereas Neon-TalinB localization was completely lost (Fig. 4C–D, “+LatA”), indicating differential dependence on the actin cytoskeleton. The partial retention of TalinA contrasts with previous reports in which both talins diffused into the cytoplasm after LatA treatment (27, 63), possibly due to differences in imaging resolution or enhanced recruitment under adhesive surface conditions. Whereas metazoan VASP localizes to both leading edges and focal adhesions (64, 65), *Dictyostelium* VASP remained restricted to the cell edge even at high WGA density (Fig. 4E). Notably, LatA treatment induced strong membrane enrichment of VASP (Fig. 4E, “+LatA”), similar to that observed for SCAR complex subunits and WASP (Fig. S5). The transmembrane adhesion proteins SibA and SadA localized to the plasma membrane and intracellular vesicles (Fig. 4F–G, “Uncoated”) and their distributions were not affected by F-actin disruption on uncoated substrates (Fig. 4F–G, “Uncoated”, +LatA). In contrast, at high WGA density, both proteins appeared as numerous foci at the ventral membrane and SibA was also strongly enriched at protrusions (Fig. 4F–G, “High WGA”). Their spatial patterns diverged after LatA treatment; SibA became enriched at the edge and central-to-posterior regions (Fig. 4F, “High WGA”, +LatA), whereas SadA was depleted from the edge (Fig. 4G, “High WGA”, +LatA). Posterior enrichment similar to that of SibA was also found for the clathrin heavy chain ChcA, which persisted even after LatA treatment (Fig. S7D).

**Figure 4.**
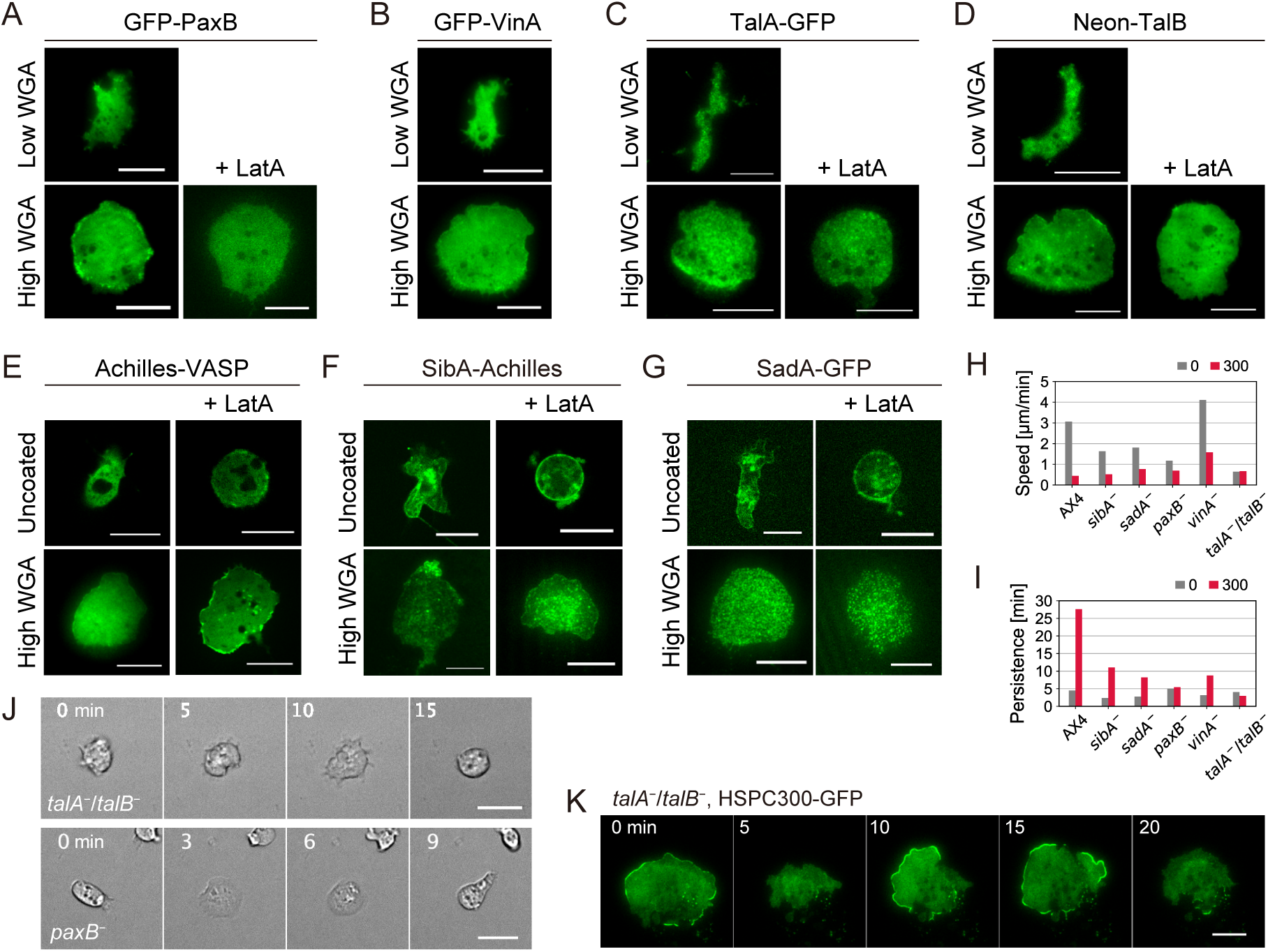
Focal adhesion-like plaques underlie sustained cell spreading and persistent migration. (**A**–**D**) TIRF images of GFP-PaxB (**A**), GFP-VinA (**B**), TalA-GFP (**C**) and Neon-TalB (**D**). Substrates were coated with 10 (“Low WGA”) and 300 µg/mL WGA (“High WGA”). (*left*) No treatment, (*right*) 3 µM LatA-treated. (**E–G**) Confocal (for “Uncoated”) and TIRF images (for “High WGA”) of Achilles-VASP (**E**), SibA-Achilles (**F**) and SadA-GFP (**G**). (*left*) No treatment, (*right*) 3 µM LatA-treated. (**H**) Migration speed and (**I**) persistence time of knockout strains. Leged, WGA concentration (µg/mL). From left to right, N = 54, 83, 57, 54, 36, 50, 64, 94, 53, 77, 53, 61 cells. (**J**) Time-lapse transmitted-light images of *talA*^−^/*talB*^−^ (*top*) and *paxB*^−^ (*bottom*) on 300 µg/mL WGA-coated substrates. Numbers, time (minutes). (**K**) Time-lapse TIRF images of HSPC300-GFP in *talA*^−^/*talB*^−^. Numbers, time (minutes) Scale bars, 10 µm in **A–G** and 20 µm in **J–K**.

The functional contributions of these adhesion molecules to migration were assessed using knockout strains. As reported previously (25, 29), *sibA*^−^ and *sadA*^−^ cells migrated at slightly reduced speeds compared with wild-type cells on uncoated substrates (Fig. 4H). On WGA-coated substrates, both mutants migrated slowly but remained highly directional, adopting a fan-shaped morphology with polarized Neon-Nap1 localization at the leading edge (Fig. 4H–I; Fig. S7E). These results indicate that SibA and SadA are dispensable for polarized migration, possibly due to functional redundancy among adhesion receptors. In contrast, *talA*^−^*/talB*^−^ cells which are unable to migrate on uncoated substrates (66) also failed to migrate on WGA-coated substrates (Fig. 4H–I). Here, *talA*^−^/*talB*^−^ cells transiently spread but retracted within minutes (Fig. 4J, *top*). These cells formed wide membrane protrusions underpinned by HSPC300-GFP (Fig. 4K, *t* = 0 and 10–15 min), followed by membrane retraction accompanied by loss of HSPC300-GFP localization (Fig. 4J, *t* = 5 and 20 min). Similarly, *paxB*^−^ cells exhibited transient spreading followed by retraction (Fig. 4J, *bottom*), and WGA coating did not enhance their migration persistence (Fig. 4I, “*paxB*^−^”). By contrast, *vinA*^−^ cells showed highly directional migration on WGA-coated substrates (Fig. 4H–I, “*vinA*^−^”). Both *paxB*^−^ and *vinA*^−^ cells were able to exhibit polarized HSPC300-GFP localization (Fig. S7F). Together, these results show that talins and PaxB are required for sustained spreading and highly persistent migration on strongly adhesive substrates.

### RacE is required for maintaining the cell rear and persistent migration

In mammalian cells, sustained spreading, which also requires talin (67), is followed by polarization through rear formation (68, 69). The persistent enrichment of myosin II at the posterior end (Fig. 5A–B) indicates stable rear formation. To examine the protrusion inducibility of the cell surface, we locally applied the chemoattractant cAMP using a microneedle positioned adjacent to migrating cells. Local stimulation with cAMP at the frontal side immediately induced membrane protrusions from the same side, and cells migrated directly toward the needle, accompanied by a transition from a fan-shaped to an elongated morphology (Fig. 5C). In contrast, stimulation from the posterior side failed to induce protrusions at the closest membrane region; instead, protrusions emerged from the lateral side, followed by cell reorientation toward the needle (Fig. 5D). This behavior resulted in a significant delay in migration toward the needle (Fig. 5E). These observations demonstrate that, even under chemoattractant induction, protrusion formation at the rear is strictly prohibited in fan-shaped cells.

**Figure 5.**
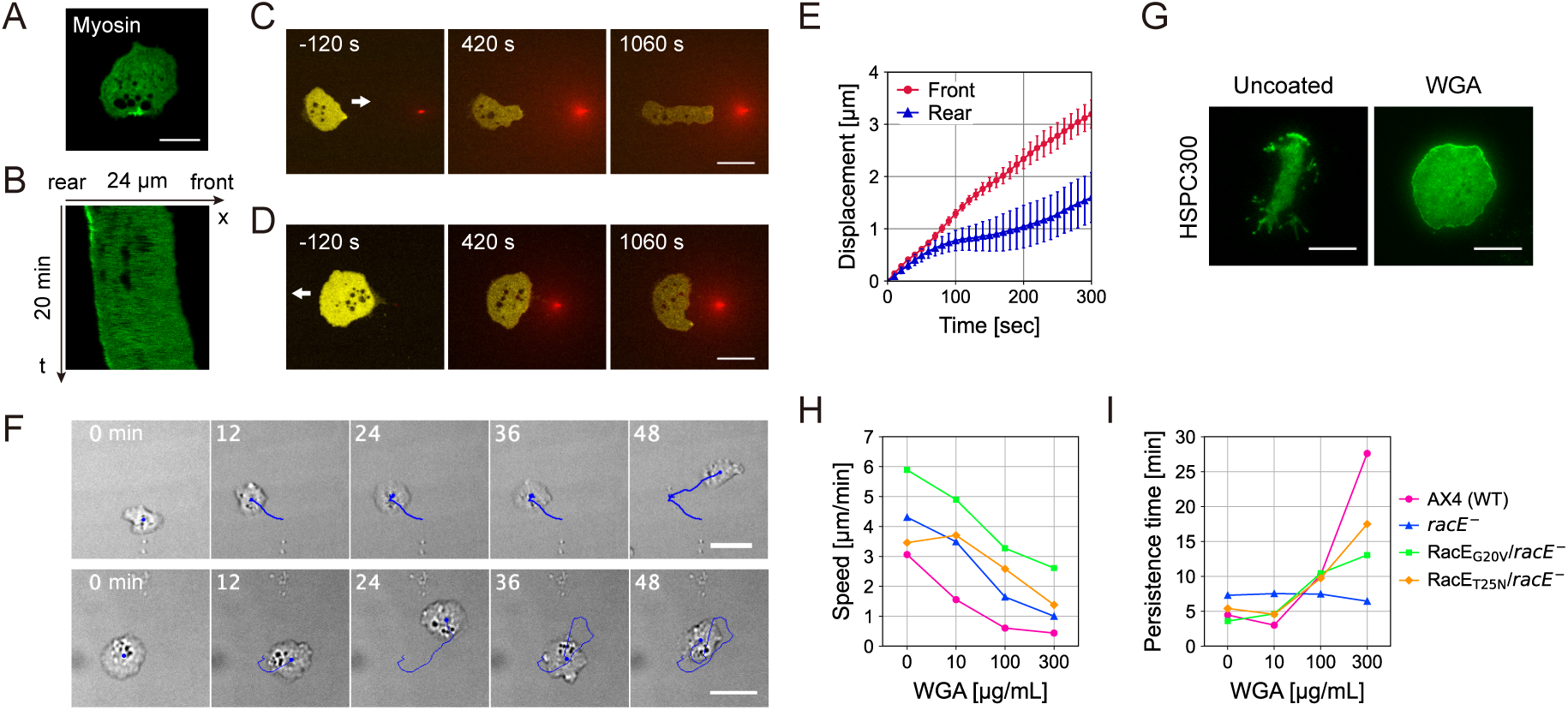
RacE is required for maintaining cell polarity and persistent migration on WGA-coated substrates. (**A**–**B**) Confocal image (**A**) and kymograph along the midline of cells (**B**) expressing GFP-MhcA on WGA-coated substrates. (**C**-**D**) Cell deformation during cAMP stimulation. A mixture of cAMP and Alexa594 (red) was applied from a microneedle to the cell front (**C**) or rear (**D**). Yellow: PH_CRAC_-mVenus. White arrows, direction of migration prior to the presentation of cAMP gradient. Substrates were coated with 300 µg/mL WGA. (**E**) Displacement of cell centroid after cAMP gradient was formed in front of (“Front”) or behind cells (“Rear”) (mean ± s.e., N = 6 and 5 cells). (**F**) *racE*^−^ cells migrated less persistently on WGA-coated substrates. Numbers, time (min). Blue line, trajectory. (**G**) TIRF images of HSPC300-GFP in *racE*^−^ cells on uncoated (*left*) and WGA-coated substrates (*right*). (**H**-**I**) Speed (**H**) and persistence time (**I**) of wild-type AX4, *racE*^−^ and *racE*^−^ expressing Neon-RacE_G20V_ or Neon-RacE_T25N_ on WGA-coated substrates. Scale bars, 10 µm in **A**, **G** and 20 µm in **C**, **D**, **F**.

RacE, the *Dictyostelium* homolog of mammalian RhoA, maintains cortical actin integrity by activating formins (70) and downstream actin crosslinkers including myosin (71). RacE-dependent cortical actin is crucial for stable cell polarization, particularly under confined spaces (70), and the cortical actin filament density is markedly reduced in *racE*^−^ cells (70). We found that when plated on WGA-coated substrates, *racE*^−^ cells exhibited abnormal migratory behaviors, such as stop-and-go movement (Fig. 5F, *top*) and frequent changes in migration direction (Fig. 5F, *bottom*). In *racE*^−^, HSPC300-GFP, which was enriched at the tips of protrusions on uncoated substrates (Fig. 5G, “Uncoated”), was broadly distributed along the entire cell edge on WGA-coated substrates (Fig. 5G, “WGA”). Consistent with these observations, the persistence time of *racE*^−^ cell trajectories did not increase on WGA-coated substrates, in contrast to wild-type cells (Fig. 5I). Expression of a constitutively active mutant RacE_G20V_ in *racE*^−^ increased both migration speed and persistence at high WGA density (Fig. 5H–I). In contrast, expression of a constitutively inactive mutant RacE_T25N_ had little effect on migration speed but unexpectedly increased migration persistence under the same conditions (Fig. 5H–I). Together, these results indicate that RacE is required for the establishment of a stable cell rear, polarized distribution of the SCAR complex, and highly persistent migration on WGA-coated substrates, in a manner that is largely independent of its canonical GTP/GDP-conversion.

### Substrate adhesion increases cell stiffness depending on talins and RacE

In metazoan cells, substrate spreading is accompanied by an increase in cell stiffness under the control of RhoA (72). To examine whether substrate adhesion similarly alters mechanical properties of *Dictyostelium* cells, we measured cell stiffness by atomic force microscopy (AFM) indentation tests. On uncoated substrates, the Young’s modulus of starved AX2 and AX4 cells was 1.4 ± 0.1 kPa and 1.7 ± 0.1 kPa, respectively. These values increased markedly to 2.5 ± 0.2 kPa and 3.1 ± 0.5 kPa when cells were plated on 300 µg/mL WGA-coated substrates (Fig. 6A). Overexpression of GbpD, a guanine nucleotide exchange factor (GEF) for Rap1 is known to enhance cell-substrate adhesion (23). Consequently, a GbpD-overexpressor exhibited modestly increased stiffness even on uncoated substrates (2.0 ± 0.2 kPa) with further increase to 3.0 ± 0.2 kPa on WGA-coated substrates (Fig. 6B, “GbpD”), in line with the notion that adhesion-dependent stiffening is not specific to WGA. Furthermore, we found that cells grown under adherent conditions were significantly stiffer than cells cultured in suspension (Fig. 6C).

**Figure 6.**
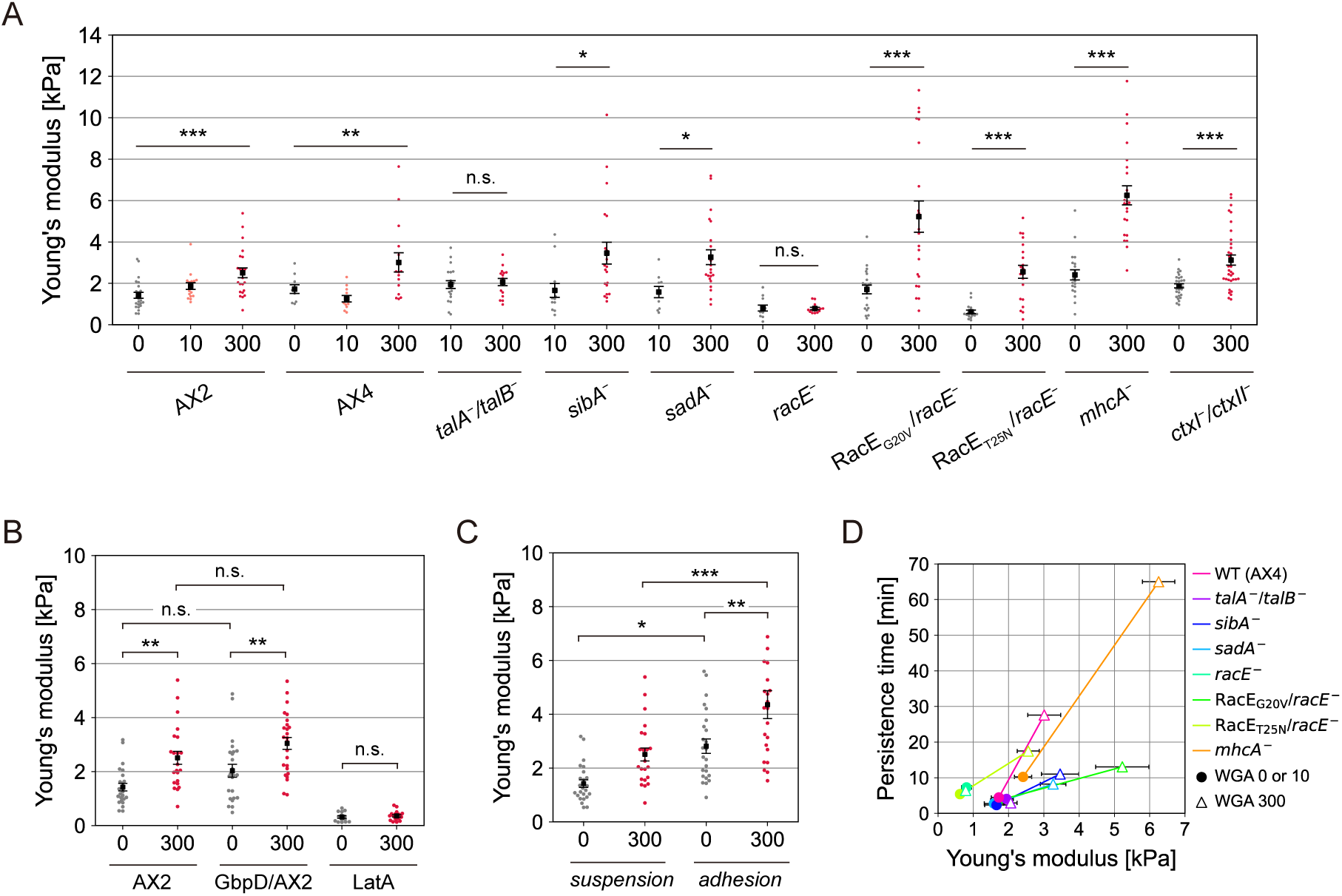
Cortical stiffness is elevated on WGA-coated substrates. (**A**) Young’s modulus of aggregation stage wild-type (AX2 and AX4) and knockout strains (*talA*^−^/*talB*^−^, *sibA*^−^, *sadA*^−^*, racE*^−^, *racE*^−^ expressing RacE_G20V_ or RacE_T25N_, *mhcA*^−^ and *ctxI*^−^/*ctxII*^−^) attached to 0, 10 and 300 µg/mL WGA-coated substrates (mean ± s.e. in black; plots for each cell; N = 9-34 cells per condition). Cells were cultured in suspension except for *mhcA*^−^ and *racE*^−^ background, which have defect in cell division in suspension. (**B**) Young’s modulus of wild-type (AX2, replicated from **A**), GbpD-overexpressing cells, and 5 µM LatA-treated AX2 cells. From left to right, N = 24, 23, 24, 26, 15, 18 cells. (**C**) Young’s modulus of AX2 cells cultured in suspension (replicated from **A**) and adherent condition. From left to right, N = 24, 23, 24 and 22 cells. (**D**) Persistence time versus Young’s modulus of wild-type and knockout strains (mean ± s.e). Plots for the same strain at low (closed circle) and high (open triangle) WGA densities are connected by solid lines. Data were compared using Student’s t-test for two groups and one-way ANOVA followed by Kramer–Tukey’s post-hoc test for more than two groups. \**p* < 0.05, \*\**p* < 0.01, and \*\*\**p* < 0.001. n.s., not significant.

Ventral adhesion structures appear to be pivotal for adhesion-induced stiffening. Cells lacking both talins (*talA*^−^*/talB*^−^) failed to increase stiffness on WGA-coated substrates, whereas *sibA*^−^ and *sadA*^−^ cells showed stiffness increases comparable to wild-type cells (Fig. 6A). Adhesion-dependent stiffening also required an intact cortical actin network. No increase in stiffness was observed in *racE*^−^ cells (Fig. 6A) or in wild-type cells treated with LatA (Fig. 6B, “LatA”). Actin crosslinking is therefore likely to underlie the increased cortical tension. However, stiffening was still observed in *racE*^−^ cells expressing either RacE_G20V_ or RacE_T25N_ (Fig. 6A, “*racE*^−^+RacE_G20V_” and “*racE*^−^+RacE_T25N_”), as well as in cells lacking myosin II heavy chain (*mhcA*^−^) or cortexillins (*ctxI*^−^/*ctxII*^−^) (Fig. 6A). These findings suggest that RacE-dependent regulation of cortical mechanics required for stiffening is largely independent of its GTP/GDP-conversion and does not rely on myosin II-mediated contractility or cortexillin-dependent crosslinking. Comparison across strains revealed a positive correlation between cortical stiffness and migratory persistence (Fig. 6D). This supports the idea that adhesion-induced cell stiffening contributes to the maintenance of front–rear polarity during fan-shaped migration.

### Substrate adhesion is required for polarization in Naegleria amoeba

Finally, we asked how substrate adhesion affects protrusion dynamics in amoeba other than *D. discoideum*. *Naegleria gruberi* is a free-living amoeba belonging to the Discoba. *Naegleria* cells possess the Arp2/3 complex and WASP family proteins (1, 2). They are highly motile and form actin-based pseudopodia (73). On uncoated borosilicate glass coverslips, *Naegleria* cells formed thin and wide lamellipodia (Fig. S8A, “Uncoated”). These expanded membranes frequently split and extended in different directions, causing the cell shape to oscillate between fan-like and dumbbell-like morphologies (73). In contrast, on substrates coated with PLL-g-PEG to block cell-substrate adhesion, cells simultaneously generated multiple narrow pseudopodia in various directions (Fig. S8A, “PEG”). These protrusions retracted within approximately 30 s, and cells failed to establish a stable front-rear polarity. Quantitative analysis revealed that protrusion lengths, measured as the distance from the cell center to the distal tip of each protrusion, were comparable between uncoated and PEG-coated substrates (Fig. S8B). In contrast, protrusion widths were significantly greater on uncoated substrates (Fig. S8C). These results indicate that substrate adhesion is dispensable for protrusion initiation but is required for the lateral expansion and stabilization of membrane protrusions, thereby enabling polarized migration of *Naegleria gruberi*.

## Discussion

In this study, we investigated how substrate adhesion reshapes cell migration, focusing primarily on the amoebozoan *Dictyostelium discoideum*. We found that *Dictyostelium* cells on strongly adhesive surfaces recapitulate key features of slow-migrating mesenchymal cells, including lamellipodia-like protrusions, focal adhesion-like plaques, and increased cortical stiffness. These features indicate that facilitated spreading on adhesive substrates is not simply a passive wetting process. Instead, adhesive surfaces trigger coordinated reorganization of the actin cytoskeleton, adhesion machinery, and contractility, resulting in a fan-shaped morphology that supports slow but highly persistent migration. Since lamellar protrusions can also form at contact-free cell surfaces, such as in migrating neutrophils (74), in macropinocytic protrusions in *Dictyostelium* and *Naegleria* cells (75), strong substrate adhesion may not be strictly required for their induction. Rather, adhesion likely acts to spatially bias membrane protrusions, guiding and stabilizing them toward persistent expansion.

While amoeboid cells are typically characterized by frequent pseudopod branching (76, 77), a feature that facilitates their rapid reorientation and exploratory migration, tight substrate adhesion increases cortical tension which likely dampens local protrusive deformations and effectively suppresses branching. Reduced branching alone, however, cannot explain persistent migration unless protrusion and retraction are spatially coordinated. The emergence of a fan-shaped cell morphology, characterized by a broad lamellipodium-like leading edge and a highly tapered trailing end, suggests that strong substrate adhesion gives rise to mechanical or signaling hysteresis required to stabilize cell polarity. Consistent with this idea, enhanced adhesion reorganizes dendritic actin networks, adhesion complexes, and Ras/Rap1 signaling into persistent front-rear gradients.

### Lamellipodia formation and mechanosensitive adhesion complexes

The SCAR complex, WASP and VASP are evolutionarily conserved activators of Arp2/3 that are essential for leading-edge formation. Enrichment of these factors at lamellipodia is a hallmark of slow-migrating metazoan cells. We showed that the broad, persistent membrane protrusions observed in *Dictyostelium* under strongly adhesive conditions were underpinned by the localization of all these factors. Enhanced cell-substrate adhesion increased the size and persistence of dendritic actin networks and promoted submembrane accumulation of Arp2/3 (Fig. S2D), SCAR (Fig. 2A), WASP (Fig. S4C), and VASP (Fig. 4E). Mutant analysis further indicated that the SCAR complex plays a predominant role over WASP in driving these lamellipodia-like protrusions. In contrast, VASP is known to be recruited to the leading edge through interaction with the SCAR complex subunit Abi (51, 78, 79). Membrane targeting of the SCAR complex is regulated by multiple factors, including interactions with acidic phospholipids (80, 81), BAR- and SH3-domain-containing proteins (82), and SCAR phosphorylation (22, 81, 83). Notably, strong enrichment of the SCAR complex at the cell edge persisted after disruption of F-actin (Fig. S5A), indicating that its membrane accumulation can be retained independently of actin polymerization. Several mechanisms may contribute to this actin-independent localization. The cell-substrate contact line may favor molecular retention through restricted diffusion, membrane curvature, or recruitment of curvature-sensing proteins. Single-molecule imaging studies have revealed that the SCAR complex is largely confined to the cell edge (84, 85), and that actin polymerization contributes to its turnover (84). Sharp membrane curvature at protrusion tips including filopodia and membrane ruffles has been proposed to influence molecular diffusion (86–88). Even in actin-free membranes, regions of high curvature can restrict propagation of Ras/PIP3 signaling domains (28, 89). Actin-independent SCAR localization may prime the membrane for deformation along the substrate, with subsequent actin polymerization amplifying protrusion growth. Adhesion-dependent expansion of membrane protrusions was likewise observed in the discoban amoeba *Naegleria gruberi* (Fig. S8) which also has WASP family proteins (1, 2). The results highlight the importance of studying their adhesion-related roles in *Naegleria*.

*Dictyostelium* cells migrating on WGA-coated substrates exhibited discontinuous, stepwise protrusion dynamics (Fig. 2E) accompanied by spatially restricted and temporally discrete cycles of PaxB assemblies (Fig. S7B–C). This behavior brings to mind the so-called stepping motility (30, 48, 52) rather than gliding motility of keratocytes and neutrophils or the traveling patch-driven migration of *Dictyostelium* under less adhesive conditions (31, 57, 59). According to a physical model, stepwise motility arises from mechanosensitive formation of spatially non-uniform adhesion complexes near the leading and trailing edges (48). Metazoan focal adhesions are stabilized by forces transmitted through F-actin and the plasma membrane, allowing them to grow near the edges (90, 91). Similarly, in *Dictyostelium*, our observations show that a strongly adhesive substrate promoted the formation of F-actin-dependent, edge-associated plaques containing PaxB (Fig. 4A). Cells lacking talins or PaxB failed to sustain spreading and polarized migration (Fig. 4H–J), resembling phenotypes of talin-depleted mouse embryonic fibroblasts (67). Taken together, persistent migration of strongly adhered *Dictyostelium* cells most likely require engagement of focal adhesion complex components at the membrane-substrate interface. In contrast to weakly adhesive conditions, where migration is talin-dependent (66) but relatively insensitive to loss of PaxB (35), our work shows that not only talins but also PaxB is indispensable for cell migration on strongly adhesive surfaces. Since SCAR and Arp2/3 activity is still enhanced in *paxB*^−^, excessive formation of branched actin may underlie the oscillatory instability of spreading dynamics (Fig. 4J) which is normally suppressed by PaxB. The differences between *paxB*^−^ and *vinA*^−^ phenotypes may reflect the distinctive domain architecture of *Dictyostelium* VinA and its apparent lack of interaction with talin (92). Likewise, the absence of VASP from ventral adhesion foci (Fig. 4E) may be related to the lack of a proline-rich domain in *Dictyostelium* vinculins, which is located in the hinge region of metazoan vinculin that mediates VASP recruitment (92–94). Taken together, sustained front-rear polarity is associated with mechanosensitive adhesion structures, which is in line with their relationship with stepping motility. In the absence of such stabilized adhesion, protrusive activity alone cannot maintain asymmetry, leading to oscillatory spreading and rapid resetting of cell polarity.

### Adhesion-dependent control of contractility and polarity

Adhesion complex formation is closely associated with increased cortical stiffness. We found that *Dictyostelium* cells grown in adherent culture were significantly stiffer than cells maintained in suspension (Fig. 6C), possibly reflecting increased F-actin content under adhesive conditions (95). In metazoan cells, substrate adhesion gives rise to a biphasic regulation of RhoA activity: an initial suppression of contractility facilitates cell spreading, followed by RhoA activation during focal adhesion maturation, leading to increased cortical tension and traction forces (72). Similar adhesion-induced stiffening has also been observed in other receptor-mediated adhesion systems (72), including cadherin-based junctions (96). Although *Dictyostelium* lacks integrins and cadherins, substrate-anchored talin can transmit forces during migration via the functional integrin homologue SibA (27, 97). We showed that substrate adhesion elevates cortical stiffness and that this regulation depends on talins, RacE, and cortical F-actin (Fig. 6A–B). This raises an interesting possibility that cell-substrate adhesion dependent regulation of cortical mechanics represents an ancestral Amorphean trait that appeared long before specialization of adhesion receptors.

The role of RacE in this process, however, is not straightforward. Both constitutively active and inactive RacE mutants restored adhesion-dependent stiffening in *racE*^−^ cells (Fig. 6A), implying that both GTP- and GDP-bound RacE contribute to cortical regulation. While this contrasts with the established role of RacE-GTP in activating formins and actin crosslinkers (70, 95), we should note that RacE-GDP has been implicated in GPCR-mediated mTORC2/Akt signaling (98). A similar apparent paradox has been reported in formin-deficient mouse fibroblasts, where adhesion-induced stiffening persists despite impaired actin polymerization (99). This led to the proposal that formins regulate cortical mechanics not only directly through actin assembly but also indirectly through focal adhesion turnover (99). Adhesion-induced cell spreading, on the other hand, can also be viewed as a form of mechanical stretching. In metazoan cells, mechanical stretching activates integrin-Src-Rap1 signaling pathway (100, 101). Similarly, in *Dictyostelium*, substrate stretching and micropipette aspiration induce myosin accumulation at the cortex (102, 103). These observations suggest that adhesion-induced stiffening may arise through multiple, partially redundant RacE-dependent mechanisms, including adhesion-mediated signaling as well as indirect effects from mechanical cell stretching.

Cell stiffening is likely related to the highly persistent cell polarity observed under strongly adhesive conditions. On such substrates, detachment at the trailing edge requires high rupture forces, thereby slowing rear retraction and increasing global cortical tension. Elevated tension would, in turn, constrain forward propagation of dendritic actin networks at the leading edge, making rear retraction rate-limiting for cell migration. Consistent with this model, *racE*^−^ cells frequently remained symmetric and quiescent or only transiently polarized on WGA-coated substrates (Fig. 5F), whereas expression of constitutively active RacE markedly increased migration speed (Fig. 5H). The symmetric morphology of *racE*^−^ cells with unusually broad distribution of the SCAR complex (Fig. 5G) resembles that of fibroblasts and keratocytes prior to symmetry breaking rear formation (68, 104). Together with its requirement for persistent migration in confined environments (70, 105), these observations suggest that RacE primarily functions to establish and maintain the global contractility required for adhesion-stabilized polarity.

While global contractility is thus the key determinant of polarization, how the cell rear is spatially determined or stabilized remains unclear. In metazoan cells, rearward actin flow and myosin accumulation reinforce front-rear polarity, rendering the rear largely passive (7, 8, 106, 107). Other modes can emerge in keratocytes, which form paired contractile regions at the rear whose mechanical balance correlates with migration directionality (108). It should be noted however that *Dictyostelium* lacks stress fibers and do not exhibit pronounced actin flows during single-cell migration. Based on our observations, one candidate mediator of the global coordination is the front-rear asymmetry of Ras and Rap1 activity (Fig. 3). The stationary gradient sharply contrasts to the actin-independent traveling waves observed in weakly adhesive conditions (52, 58, 61). The fact that the gradient also persists even after F-actin disruption indicates that cell polarity can be maintained through adhesion-dependent but actin-independent mechanisms. Classical adhesion complexes per se are unlikely to serve this purpose, because disruption of F-actin abolished the polarized distribution of both TalB- and PaxB-enriched foci (Fig. 4A, D). While TalA-enriched foci were still visible under LatA treatment, their spatial distribution showed no clear asymmetry (Fig. 4C). Of the proteins examined, SibA and ChcA displayed front-rear asymmetry which inversely correlated with that of Rap1-GTP even after LatA treatment (Fig. 4F; Fig. S7D). Recent studies have increasingly characterized clathrin lattices as a distinct class of adhesomes in metazoa that work independently of the actomyosin-linked integrin machineries (109); however, their relationship to Rap1 remains unknown. In *Dictyostelium*, although Rap1-GTP directly binds to TalB and not to TalA, constitutive Rap1 activation enhances adhesion in *talB*^−^ cells (41), suggesting that its effect on adhesion extends beyond this specific interaction. Among recently characterized proteins that interact with Rap1, IqgC localization depends on PaxB (44) and consequently on actin, whereas the Mst1 homolog KrsB (43) has yet to be characterized regarding its subcellular distribution. Recent studies have shown that Rap1 is essential for B cell migration (110) and regulates integrin-independent T cell polarization through Mst1 (111). Our findings point to two complementary modes of polarity control: one that is actin-dependent and mechanically coordinated, and the other that is adhesion-dependent yet actin-independent. Taken together, our observations may reflect an evolutionarily conserved dual-layered architecture of polarity regulation.

## Conclusion

Strong substrate adhesion induces a mesenchymal-like, fan-shaped migration of *Dictyostelium* cells. It is accompanied by the persistent localization of the SCAR complex, focal adhesion-like plaques, persistent front-rear gradients of Ras/Rap and SibA, and increased cortical stiffness. While F-actin drives protrusive and contractile forces and supports ventral adhesion foci, the front-rear polarity of Ras/Rap activity is maintained independently of F-actin and classical focal adhesion complex components PaxB. Our results show that *Dictyostelium* cells can adopt a persistently polarized state through adhesion-dependent mechano-regulation, suggesting an evolutionary ancient coupling between cell-substrate adhesion, contractility, and cell polarity.

## Materials and Methods

### Cell preparation and live-cell imaging

All the plasmids and cell strains used in this study are listed in Table. S1–2. Axenic strains of *Dictyostelium discoideum* were cultured as previously described (112). For live-cell imaging, cells were washed twice, resuspended in developmental buffer (DB) at 5×10^6^ cells/mL and shaken at 22°C, 155 rpm for 1 h. Cells were then pulsed with cAMP at final concentration of 50 nM every 6 min for 4.5 h. Starved cells were collected by centrifugation, resuspended in DB, and plated on glass coverslips at 3×10^3^ cells/cm^2^ for at least 1 h. Images were acquired using an inverted microscope (IX83 or IX81, Olympus) equipped with a laser confocal scanning unit (CSU-W1 or CSU-X1, YOKOGAWA) and an EMCCD camera. To observe cell deformation under a spatial gradient of cAMP concentration, 100 nM cAMP and 10 µg/mL Alexa594 were loaded into Femtotips II (Eppendorf) and pressurized at 100 hPa. All live-cell imaging was performed at 22°C. *Naegleria gruberi* strain NEG-M was cultured as previously described (73). Cells were grown and observed in HL5 medium at 30°C.

### Substrate modification

To promote cell-substrate attachment, wheat germ agglutinin lectin (WGA; Wako, 126-02811), poly-L-lysine (PLL; Sigma, P4832) and Cell-tak (CTK; Wako, 646-54971) were used. WGA was dissolved in milliQ water at 1 mg/mL and stored in aliquots at −30°C. PLL and CTK were stored at 4°C. Borosilicate glass coverslips (Matsunami, C018001, φ25 mm) were cleaned by sequential sonication in alkaline solution and ethanol, and then attached to the bottom of 35-mm culture dishes as previously described (113). Prior to coating, dishes were treated with air plasma (Harrick Plasma, PDC-32G) at 400 mTorr for 2 min to improve wettability. For WGA coating, the surface was covered with 100 µL of WGA solution (0-300 µg/mL in DB) and incubated for 1 h at room temperature (RT), followed by rinsing with DB. For PLL coating, the surface was covered with 100 µL of PLL solution (0-100 µg/mL in milliQ water) and incubated for 1 h at RT, after which the surface was rinsed with milliQ water and air-dried. For CTK coating, the surface was covered with 20 µL of CTK solution (0-300 µg/mL in PBS) and allowed to dry at RT. To inhibit cell adhesion, PLL(20)-g[3,5]-PEG(2) (PLL-g-PEG; SuSoS AG) was dissolved in milliQ water at 1 mg/mL and stored in aliquots at −30°C. Plasma-treated coverslips were covered with 100 µL of 100 µg/mL PLL-g-PEG solution for 1 h, rinsed thoroughly with water, and air-dried.

### AFM measurements

Starved *Dictyostelium* cells prepared as described above were plated on WGA-coated coverslips 1 h before measurements. Force spectroscopy was performed using a NanoWizard 3 Ultra AFM (Bruker) mounted on an inverted microscope (IX70, Olympus). The entire system was installed on an active vibration isolation table (Hertz). A PNP-TR cantilever (four-sided pyramidal tip, Nanoworld) was used. The nominal spring constant was 0.32 N/m and was calibrated for each experiment using the thermal noise method. The photodetector sensitivity was determined by linear fitting of the slope of the force-distance curve acquired on a glass substrate. Indentation tests were repeated more than 16 times per cell, with approach and retraction speeds of 2 µm/s. The elastic modulus of cells attached to the glass was determined by fitting the Hertz model to the force-indentation curves using an indentation depth < 1 µm. All measurements were performed at 22°C.

### Cell shape and trajectory analysis

Image analysis was performed using ImageJ, Python, and Microsoft Excel. To quantify cell shape and migration dynamics, transmitted-light and confocal images of GFP-Lifeact expressing cells were acquired at 1-min intervals for 50 min using a 20× objective. Fluorescence images were used to generate cell masks, from which cell area and the lengths of the major and minor axes were extracted at each time point by ellipse fitting. Cell trajectories were obtained by automatic or manual tracking of cell centroids and were smoothed by averaging three consecutive positions. The direction of cell migration was determined from the displacement vector between consecutive frames and was defined as the front-back (FB) axis of the cell. The major or minor axis whose orientation most closely matched the FB direction was assigned as the FB axis, while the orthogonal axis was defined as the left-right (LR) axis. Cell area and the ratio of front-back length to left-right width (FB/LR) were averaged for each cell trajectory. The mean square displacement (MSD) of cell trajectories was calculated, and cell speed and persistence time were obtained by fitting the data to a persistent random walk model. Trajectory persistence was also independently quantified for individual cells as the ratio of net displacement to total path length.

### Boundary tracking analysis

To quantify the spatiotemporal dynamics of the SCAR complex and membrane protrusion, boundary tracking analysis was performed (47). Briefly, cell masks were generated from fluorescence images, and cell contours were extracted. Each contour was divided into 100 segments, and the centroid coordinates of each segment were determined. To establish correpondence between segments at successive time points, distances between each segment at time point *t* and those at *t* + 1 were calculated, and segments were linked so as to minimize the sum of squared distances. The normal component of the resulting displacement was defined as the local protrusion speed. The fluorescence intensity of HSPC300-GFP for each segment was calculated as the average intensity within a region extending 5 pixels (approximately 340 nm) inward from the cell boundary and normalized by the average intensity of a region located an additional 10 pixels inward, averaged over the entire contour. Protrusion speed and fluorescence intensity for each segment and time point were visualized as heatmaps (Fig. 2B–E) and 2D histograms (Fig. S3F–G). Using the resulting 2D matrices, spatiotemporal autocorrelations were calculated. Cross-sections at zero segment shift (Δ*x* = 0) and zero time lag (*τ* = 0) were plotted as temporal and spatial autocorrelation functions, respectively (Fig. 2F–I).

### Quantification of front-rear fluorescence gradients along the cell axis

To evaluate front-rear gradients of activated small GTPases and PIP3 in cells attached to WGA-coated substrates (Fig. 3C), timelapse TIRF images were acquired at 5-s intervals for intact cells and 1-min intervals for LatA-treated cells. Kymographs were taken along the cell midlines with a width of 10 pixels. Fluorescence intensity at each time point was normalized to the maximum intensity within the same frame to correct for photobleaching. Cell edges were identified as positions of maximal and minimal spatial derivatives of the fluorescence intensity profiles. Using the detected front and rear edges, fluorescence profiles were spatially aligned and rescaled to a normalized cell length coordinate, with the front and rear edges corresponding to 0 and 1, respectively. The normalized cell length was divided into 21 bins, and fluorescence intensity was averaged within each bin. For each cell, time-averaged spatial profiles were calculated and subsequently averaged across cells.

## Acknowledgements

The authors thank present and past members of the Sawai lab for various technical and scientific inputs. We thank Jean-Paul Rieu for introducing us to WGA and for insightful discussions, Thomas Böddeker for beneficial discussion, Robert Insall and Dicty Stock Center (DSC) for pDM459 HSPC300-GFP and *pirA*^−^, Tatsuo Kinashi for pEGFP-N1/RalGDS-RBD, Tobias Meyer and Addgene for PLCdelta-PH (Addgene #21179), Igor Weber for GFP-CI, James Spudich and DSC for pBIG-GFP-myo, Taro Uyeda, Akira Nagasaki and the National BioResource Project (NBRP) Nenkin for pBIG GFP-VinA, pBIG GFP-PaxB, *paxB*^−^, *vinA*^−^, Masatsune Tsujioka and NBRP for pTA15 TalA-GFP and *talA*^−^/*talB*^−^, Petra Fey and DSC for *sadA*^−^ cells expressing SadA-GFP, Robert Kay and NBRP for *pi3k1-5*^−^, Peter Devreotes and DSC for *pten*^−^, Miho Iijima and Hiroshi Senoo for pTXGFP-LactC2 and *racE*^−^, Günther Gerisch and NBRP for *ctxI*^−^/*ctxII*^−^ and Douglas Robinson and DSC for *mhcA*^−^. This work was supported by grants from Japan Science and Technology Agency (JST) CREST JPMJCR1923, Japan Society for Promotion of Science (JSPS) KAKENHI JP25K18459 (to G.H.), JP19H05801, JP25H01363 (to S.S.), research grant from the Mitsubishi Foundation (to S.S.), and in part by Joint Research by Exploratory Research Center on Life and Living Systems (ExCELLS) Grant 18-204 (to S.S.), JSPS KAKENHI JP17H01812, JP17H05992, JP18H04759, JP19H05416, JP23H00384, JP25H01771 and JP25K22490, JP26H00950 (to S.S.).

## Data Availability

All data included in this paper and SI Appendix will be available at RIKEN Systems Science Biological Dynamics database repository.

