## Supplementary file for "Adhesion-mediated transition to a mesenchymal-like, fan-shaped migration mode in *Dictyostelium discoideum*"

*Corresponding author: Satoshi Sawai

**This PDF file includes:**

Table S1 to S2

Figures S1 to S8

SI References

**Table S1. Plasmid list used in this study**

| # | Plasmid name | Backbone | Reference |
|---|---|---|---|
| 1 | pDM304 GFP-Lifeact | pDM304 | (1) |
| 2 | pDM1208 mCherry-Lifeact | pDM1208 | This study |
| 3 | pDM459 HSPC300-GFP | pDM459 | (2) |
| 4 | pDM304 Neon-Nap1 | pDM304 | This study |
| 5 | pDM304 PIR121-Achilles | pDM304 | This study |
| 6 | pDM1210 PIR121-mCherry | pDM1210 | This study |
| 7 | pDM304 mVenus-WASP-A | pDM304 | This study |
| 8 | pDM304 Achilles-WASP-A | pDM304 | This study |
| 9 | pDM358 IBARa-mScarlet-I | pDM358 | This study |
| 10 | pDM304 PH$_{CRAC}$-mVenus | pDM304 | This study |
| 11 | pDM1209 PH$_{Akt}$-GFP | pDM1209 | This study |
| 12 | pDGFP-CI | pDGFP-MCS | (3) |
| 13 | pHyg PH$_{PLC\delta}$-EGFP | pHyg | This study |
| 14 | pTX GFP-LactC2 | pTX | (4) |
| 15 | pDM323 CRIB$_{PakB}$-GFP | pDM323 | This study |
| 16 | pHyg RBD$_{RalGDS}$-GFP | pHyg | This study |
| 17 | pA15 TalinA-GFP | pA15 | (5) NBRP Nenkin (ID: G90576) |
| 18 | pDM304 Neon-TalinB | pDM304 | This study |
| 19 | pBIG GFP-VinA | pBIG | (6) NBRP ID: G90300 |
| 20 | pBIG GFP-PaxB | pBIG | (6) NBRP ID: G90301 |
| 21 | pDM326 SibA-Achilles | pDM326 | This study |
| 22 | pDM326 Achilles-VASP | pDM326 | This study |
| 23 | pBIG GFP-MhcA | pBIG | (7) DSC ID: 381 |
| 24 | pDM304 Neon-RacE$_{G20V}$ | pDM304 | This study |
| 25 | pDM304 Neon-RacE$_{T25N}$ | pDM304 | This study |
| 26 | pDM358 mScarlet-I-GbpD | pDM358 | This study |

**Table S2. Cell strain list used in this study**

| # | Strain name | Backbone | Reference |
|---|---|---|---|
| 1 | AX4 (*Dictyostelium discoideum*) | | Lab stock |
| 2 | AX2 (*Dictyostelium discoideum*) | | Lab stock |
| 3 | NEG-M (*Naegleria gruberi*) | | Lab stock |
| 4 | *pirA*$^{-}$ | AX3 | (8) DSC ID: DBS0236780 |
| 5 | *wasA*$^{-}$ | AX2 | This study |
| 6 | *sibA*$^{-}$ | AX4 | This study |
| 7 | *sadA*$^{-}$ | AX4 | This study |
| 8 | *pi3k1-5*$^{-}$ | AX2 | (9) NBRP ID: S00405 |
| 9 | *pten*$^{-}$ | AX2 | (10) DSC ID: DBS0236830 |
| 10 | *paxB*$^{-}$ | AX2 | (6) NBRP ID: S00317 |
| 11 | *vinA*$^{-}$ | AX2 | (6) NBRP ID: S00551 |
| 12 | *talA*$^{-}$/*talB*$^{-}$ | AX2 | (5) NBRP ID: S00028 |
| 13 | *racE*$^{-}$ | AX2 | (11) DSC ID: DBS0350271 |
| 14 | *mhcA*$^{-}$ | JH10 | (12) DSC ID: DBS0236379 |
| 15 | *ctxI*$^{-}$/*ctxII*$^{-}$ | AX2 | (13) NBRP ID: S00404 |
| 16 | GFP-Lifeact/AX4 | AX4 | (14) |
| 17 | HSPC300-GFP/AX4 | AX4 | This study |
| 18 | HSPC300-GFP/Lifeact-mRFPmars/AX4 | AX4 | (1) |
| 19 | GFP-Arp2/AX4 | AX4 | (1) |
| 20 | HSPC300-GFP/AX2 | AX2 | This study |
| 21 | HSPC300-GFP/*wasA*$^{-}$ | *wasA*$^{-}$ | This study |
| 22 | HSPC300-GFP/*pirA*$^{-}$ | *pirA*$^{-}$ | This study |
| 23 | PIR121-mCherry/*pirA*$^{-}$ | *pirA*$^{-}$ | This study |
| 24 | mVenus-WASP-A/AX4 | AX4 | This study |

| | | | |
|---|---|---|---|
| 25 | Achilles-WASP-A/AX4 | AX4 | This study |
| 26 | mVenus-WASP-A/*pirA*$^{-}$ | *pirA*$^{-}$ | This study |
| 27 | mVenus-WASP/PIR121-mCherry/*pirA*$^{-}$ | *pirA*$^{-}$ | This study |
| 28 | Neon-Nap1/AX4 | AX4 | This study |
| 29 | Neon-Nap1/*pirA*$^{-}$ | *pirA*$^{-}$ | This study |
| 30 | Neon-Nap1/PIR121-mCherry/*pirA*$^{-}$ | *pirA*$^{-}$ | This study |
| 31 | mScarlet-I-IBARa/AX4 | AX4 | This study |
| 32 | $PH_{CRAC}$-GFP/AX4 | AX4 | (14) |
| 33 | $PH_{CRAC}$-mVenus/AX4 | AX4 | This study |
| 34 | $PH_{AKT}$-GFP/AX4 | AX4 | This study |
| 35 | $PH_{CRAC}$-RFP/PTEN-GFP/AX4 | AX4 | (14) |
| 36 | GFP-Cortexillin I/Lifeact-mRFPmars/*ctxI*$^{-}$ | *ctxI*$^{-}$ | (14) |
| 37 | $PH_{PLC\delta}$-EGFP/AX4 | AX4 | This study |
| 38 | GFP-LactC2/AX4 | AX4 | This study |
| 39 | HSPC300-GFP/*pi3k1-5*$^{-}$ | *pi3k1-5*$^{-}$ | This study |
| 40 | HSPC300-GFP/*pten*$^{-}$ | *pten*$^{-}$ | This study |
| 41 | GFP-$RBD_{Raf1}$/AX2 | AX2 | (14) |
| 42 | $RBD_{RalGDS}$-GFP/Lifeact-mRFPmars/AX4 | AX4 | This study |
| 43 | $CRIB_{PakB}$-GFP/AX4 | AX4 | This study |
| 44 | $PH_{Akt}$-GFP/mCherry-Lifeact/AX4 | AX4 | This study |
| 45 | GFP-PaxB/AX4 | AX4 | This study |
| 46 | GFP-VinA/AX4 | AX4 | This study |
| 47 | TalinA-GFP/AX4 | AX4 | This study |
| 48 | Neon-TalinB/AX4 | AX4 | This study |
| 49 | Achilles-VASP/AX4 | AX4 | This study |
| 50 | SibA-Achilles/AX2 | AX2 | This study |
| 51 | SadA-GFP/*sadA*$^{-}$ | *sadA*$^{-}$ (AX3) | (15)<br>DSC ID:<br>DBS0236922 |
| 52 | Neon-Nap1/*sibA*$^{-}$ | *sibA*$^{-}$ (AX4) | This study |
| 53 | Neon-Nap1/*sadA*$^{-}$ | *sadA*$^{-}$ (AX4) | This study |
| 54 | HSPC300-GFP/*paxB*$^{-}$ | *paxB*$^{-}$ | This study |

| | | | |
|---|---|---|---|
| 55 | HSPC300-GFP/*vinA*$^{-}$ | *vinA*$^{-}$ | This study |
| 56 | HSPC300-GFP/*talA*$^{-}$/*talB*$^{-}$ | *talA*$^{-}$/*talB*$^{-}$ | This study |
| 57 | GFP-Myo/AX4 | AX4 | (14) |
| 58 | HSPC300-GFP/*racE*$^{-}$ | *racE*$^{-}$ | This study |
| 59 | Neon-RacE$_{G20V}$/*racE*$^{-}$ | *racE*$^{-}$ | This study |
| 60 | Neon-RacE$_{T25N}$/*racE*$^{-}$ | *racE*$^{-}$ | This study |
| 61 | mScarlet-I-GbpD/AX4 | AX4 | This study |

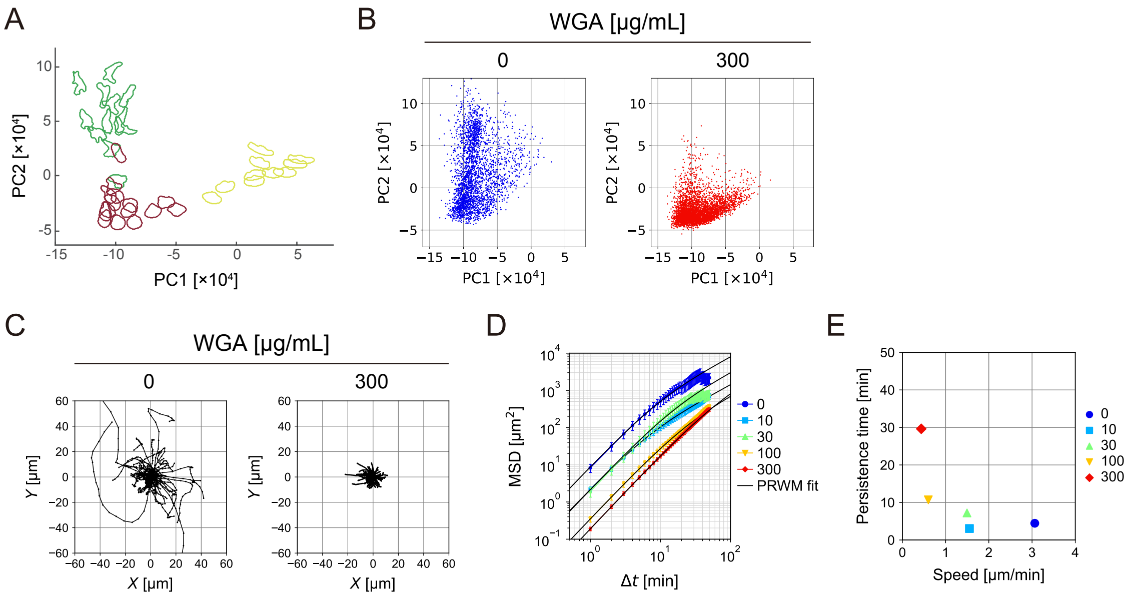


**Figure S1. Projection of cell morphologies onto a PCA-defined shape space and MSD analysis**

(**A**) Deep learning-based classification of diverse cell morphologies. Cell shapes are mapped into a principal component analysis (PCA) space. Shown are reference morphologies; green, starved *Dictyostelium*, red: neutrophil-like HL60, yellow: fish keratocytes. (**B**) *Dictyostelium* cell morphologies on uncoated (*left,* N = 2950 plots) and WGA-coated (*right,* , N = 4300 plots) substrates projected into the PCA space. Plots represent all cells across time frames. (**C**) Cell trajectories of aggregation-stage AX4 cells on uncoated (*left*) and WGA-coated substrates (*right*) over a 30 min period. (**D**) Mean square displacement (MSD) of cell trajectories. Legend, WGA concentration (μg/mL). N = 54, 74, 78, 62, 83 cells. Solid lines, fitting curves based on the persistent random walk model (PRWM). (**E**) Scatter plot of speed versus persistence time obtained from curve-fitting to MSD curves. Legend, WGA concentration (μg/mL).

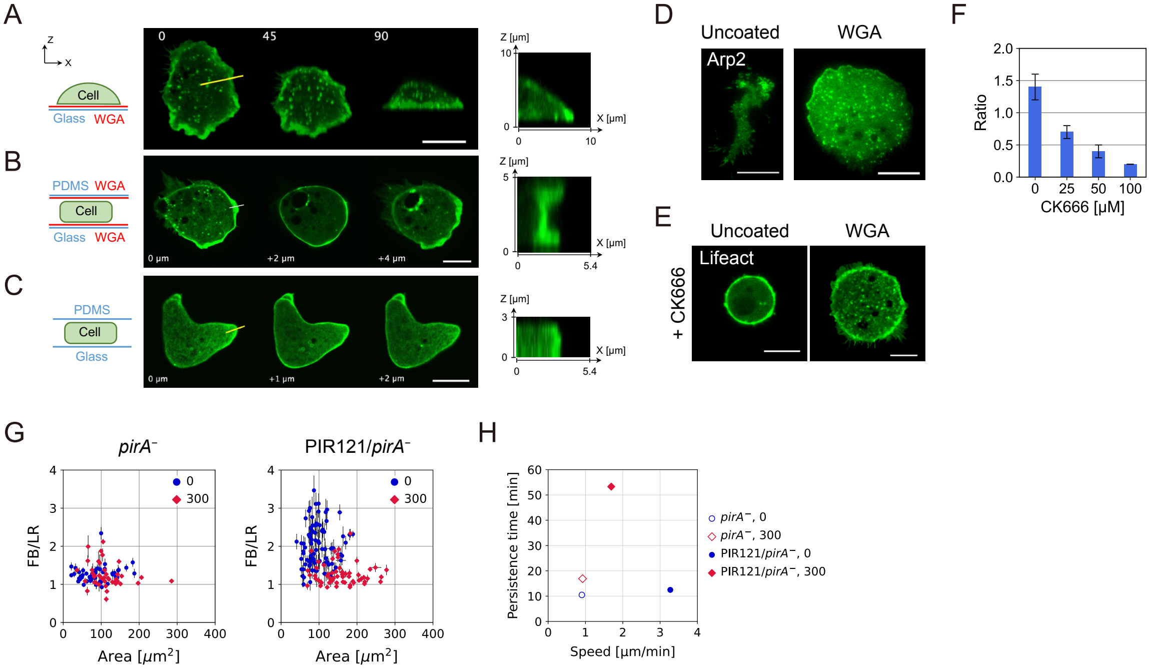


**Figure S2. Fan-shaped, persistent migration depends on the SCAR complex and Arp2/3-mediated actin nucleation.**

(**A**) Maximum intensity projections of Zstacks (10 μm thickness, 0.5 μm steps) of a GFP-Lifeact-expressing cell on the WGA-coated substrate, viewed at 0°, 45°, and 90°. (*right*) XZ cross-section along the yellow line in the center image. (**B**) Confocal images of a GFP-Lifeact-expressing cell placed between WGA-coated glass and PDMS substrates. Shown are confocal sections near the glass surface (z = 0 μm), mid-plane (z = 2 μm), and near the PDMS surface (z = 4 μm). (*right*) corresponding XZ cross-section. (**C**) Confocal images of a GFP-Lifeact-expressing cell placed between uncoated glass and PDMS substrates. Confocal sections are shown at z = 0, 1, and 2 μm. (*right*) corresponding XZ cross-section. (**D**) TIRF images of GFP-Arp2 in cells attached to uncoated (*left*) and WGA-coated substrates (*right*). (**E**) Confocal images of GFP-Lifeact in cells treated with 100 μM CK666. Scale bars, 10 μm. (**F**) Ratio of cell speed after CK666 treatment to that before treatment (mean ± s.e., N = 8, 8, 7, 7 cells). (**G**) Scatter plot of cell shape features; the spreading area (Area) and the ratio of front-back length to left-right width (FB/LR). *pirA*$^{-}$ (*left,* N = 45 and 42 cells) and *pirA*$^{-}$ expressing PIR121-Achilles (*right,* N = 90 and 60 cells). Legend, WGA concentration (μg/mL). (**H**) Scatter plot of speed versus persistence time.

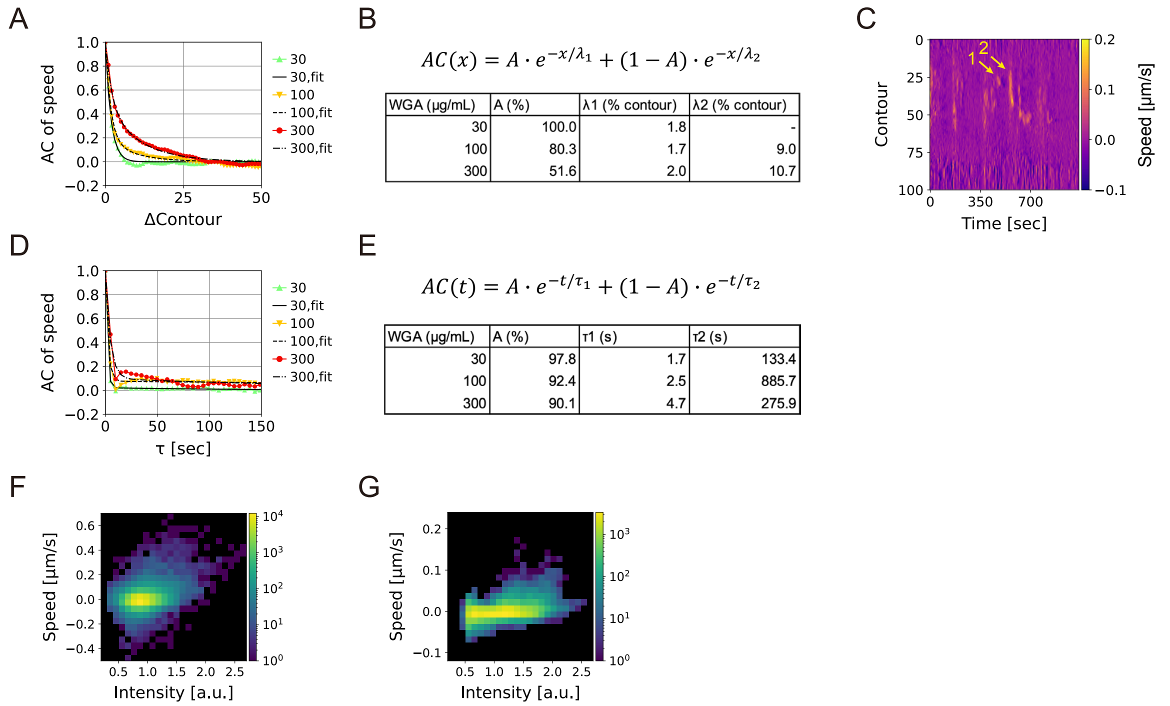


$$AC(x) = A \cdot e^{-x/\lambda_1} + (1 - A) \cdot e^{-x/\lambda_2}$$

| WGA (μg/mL) | A (%) | λ1 (% contour) | λ2 (% contour) |
|---|---|---|---|
| 30 | 100.0 | 1.8 | - |
| 100 | 80.3 | 1.7 | 9.0 |
| 300 | 51.6 | 2.0 | 10.7 |

$$AC(t) = A \cdot e^{-t/\tau_1} + (1 - A) \cdot e^{-t/\tau_2}$$

| WGA (μg/mL) | A (%) | τ1 (s) | τ2 (s) |
|---|---|---|---|
| 30 | 97.8 | 1.7 | 133.4 |
| 100 | 92.4 | 2.5 | 885.7 |
| 300 | 90.1 | 4.7 | 275.9 |

**Figure S3. Fan-shaped migration on WGA-coated substrates depends on dendritic actin networks**

(**A**, **B**, **D**, **E**) Autocorrelation of protrusion speed in space (**A**) and time (**D**). Legend, WGA concentration (μg/mL). N = 4 cells each. Curves were fitted with biexponential functions shown in **B** and **E**. (**B**, **E**, *bottom*) Tables of fitting parameters. (**C**) Heatmap showing spatiotemporal distribution of local protrusion speed along the cell boundary on 300 μg/mL WGA-coated substrates. Yellow arrows indicate narrow ("1") and wide ("2") protrusions. (**F**–**G**) 2D histograms of local HSPC300-GFP intensity and protrusion speed. Counts are shown in pseudocolor. Cells on WGA-coated substrates at (**F**) 30 (N = 5 cells) and (**G**) 300 μg/mL (N = 4 cells).

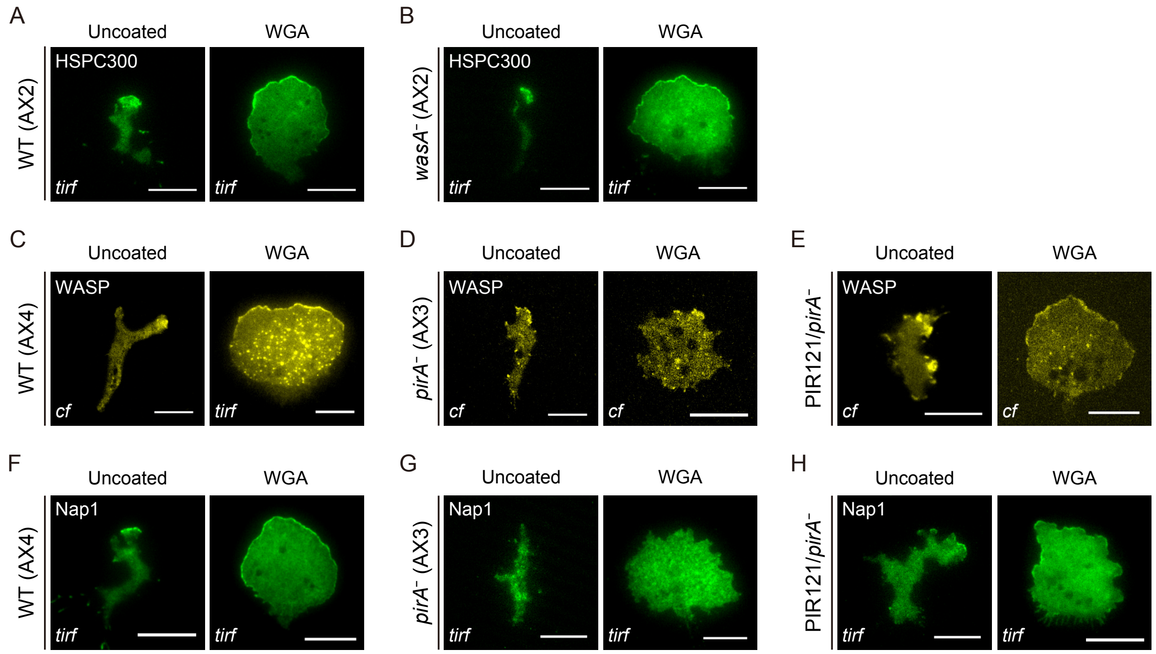


**Figure S4. WASP is recruited to the leading edge independently of the SCAR complex and cell-substrate adhesion**

TIRF and confocal images (indicated with "*tirf*" or "*cf*" at the bottom left of each image) of fluorescent protein-tagged WASP and the SCAR complex subunits on uncoated and WGA-coated substrates. (**A–B**) HSPC300-GFP in AX2 and *wasA*⁻. (**C–H**) Achilles-WASP-A (**C**) and Neon-Nap1 (**F**) in AX4. mVenus-WASP-A (**D**–**E**) and Neon-Nap1 (**G**–**H**) in *pirA*⁻, and *pirA*⁻ expressing PIR121-mCherry. Scale bars, 10 μm.

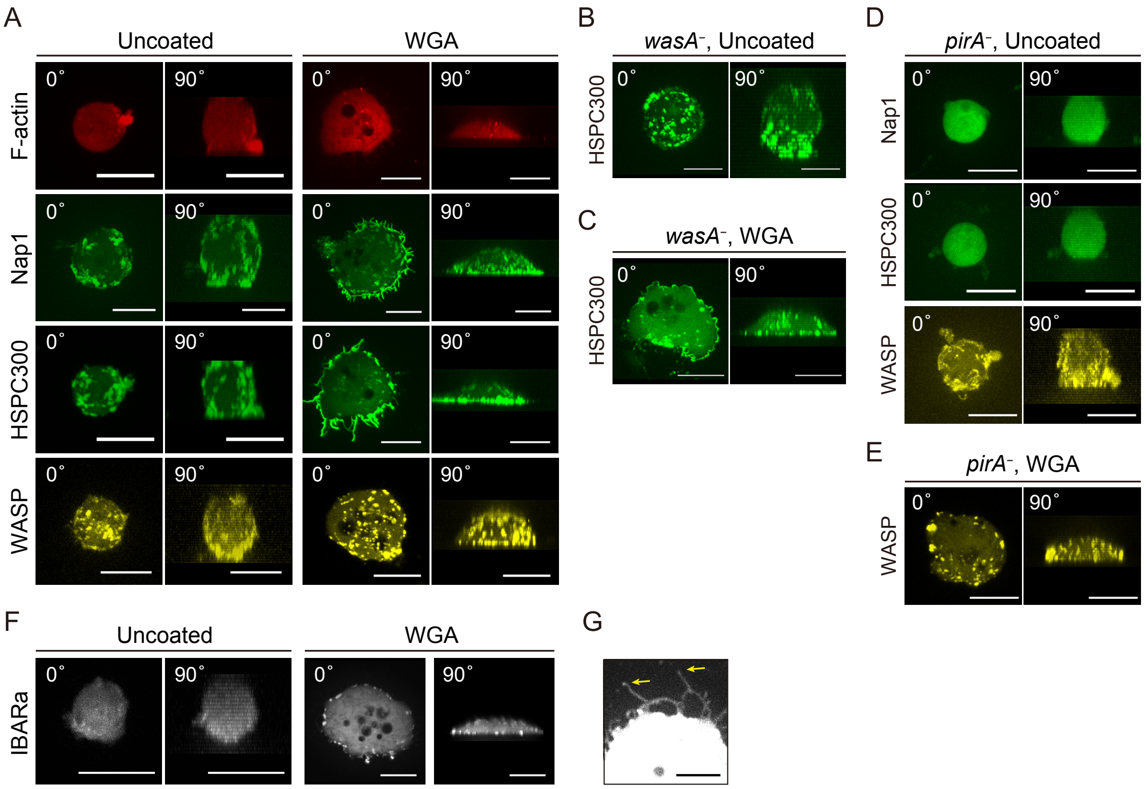


**Figure S5. F-actin disruption induces strong condensation of the SCAR complex, WASP and I-BAR protein**

(**A**) Maximum intensity projection of confocal images of Lifeact-mRFPmars, Neon-Nap1, HSPC300-GFP and mVenus-WASP-A in cells treated with 3 µM LatA on uncoated (*left*) and WGA-coated (*right*) substrates. Zstacks were taken at 0.5 µm intervals and projected onto the XY and XZ planes (indicated at the top left of each image as "0°" and "90°", respectively). (**B**–**C**) HSPC300-GFP in LatA-treated *wasA*$^-$ on (**B**) uncoated and (**C**) WGA-coated substrates. (**D**) Neon-Nap1, HSPC300-GFP and mVenus-WASPA in LatA-treated *pirA*$^-$ on uncoated substrates. (**E**) mVenus-WASPA in LatA-treated *pirA*$^-$ on WGA-coated substrates. (**F**) IBARa-mScarlet-I in LatA-treated AX4 cells on uncoated (*left*) and WGA-coated (*right*) substrates. (**G**) Confocal image of IBARa-mScarlet-I localized at the tips of filamentous membrane extensions on WGA-coated substrates. Scale bars, 10 µm in **A**–**F** and 5 µm in **G**.

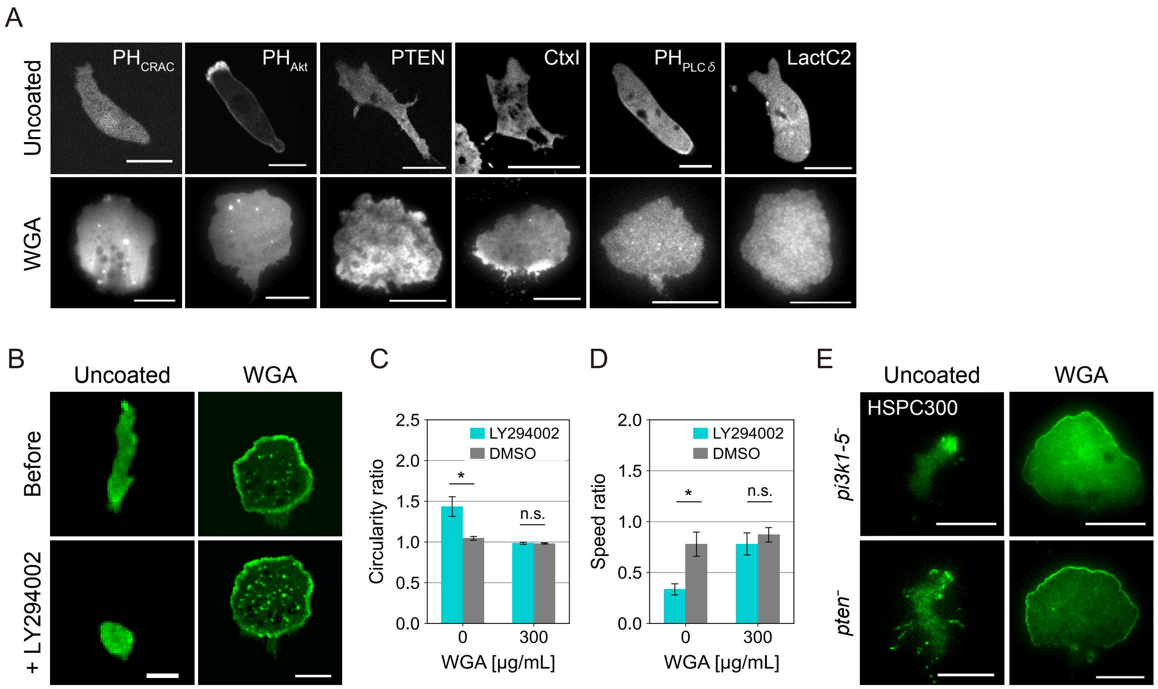


**Figure S6. Fan-shaped migration on WGA-coated substrates is independent of PIP3**

(**A**) Confocal (*top*, uncoated) and TIRF images (*bottom*, WGA-coated) of cells expressing $PH_{CRAC}$-GFP, $PH_{Akt}$-GFP, PTEN-GFP, GFP-CtxI, $PH_{PLC\delta}$-EGFP and GFP-LactC2. (**B**) Confocal images of GFP-Lifeact-expressing cells before (*top*) and after (*bottom*) treatment with 50 μM LY294002 on uncoated (*left*) and WGA-coated (*right*) substrates. (**C–D**) Ratio of circularity (**C**) and speed (**D**) of cells treated with LY294002 and DMSO to pretreatment values (mean ± s.e., N = 6, 26, 22 and 17 cells from left to right). Welch's t-test. *$p < 0.05$, n.s., not significant. (**E**) TIRF images of HSPC300-GFP in *pi3k1-5*$^-$ (*top*) and *pten*$^-$ (*bottom*) on uncoated (*left*) and WGA-coated (*right*) substrates. Scale bars, 10 μm.

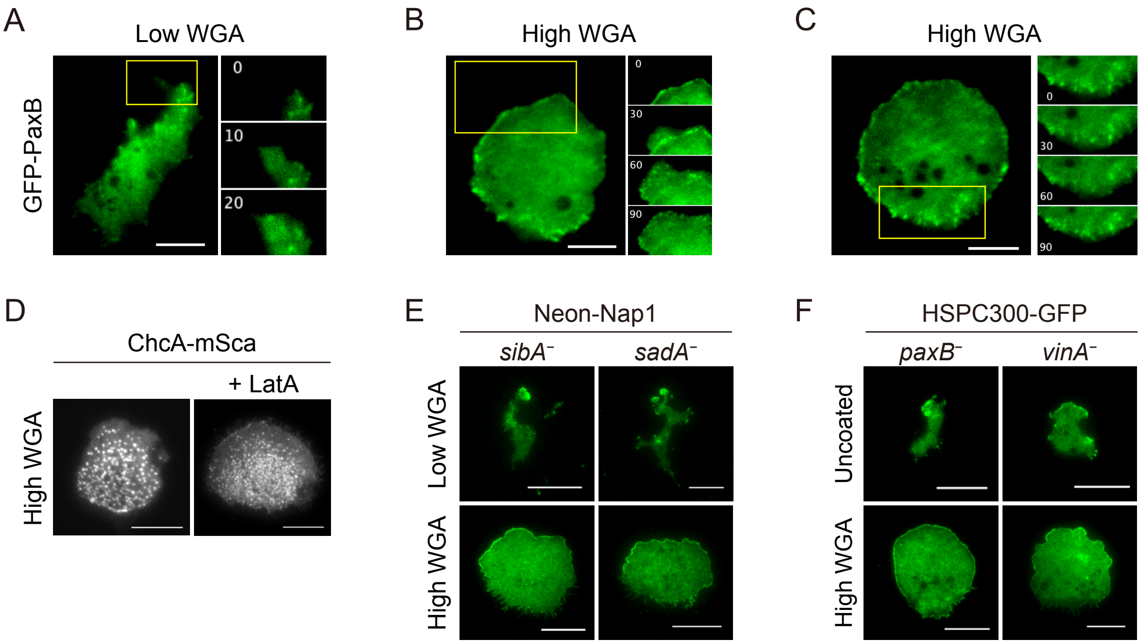


**Figure S7. Polarized SCAR subunit localizations in adhesion protein knockout strains**

(**A–C**) TIRF images of GFP-PaxB in AX2 on WGA-coated substrates (**A**, 30 µg/mL; **B** and **C,** 300 µg/mL). (inset) Timelapse images at the region indicated by yellow box in *left*. Numbers, time (s). (**D**) TIRF images of ChcA-mSca in intact (*left*) and LatA-treated (*right*) AX4 on WGA-coated substrates. (**E**) TIRF images of Neon-Nap1 in *sibA*$^{-}$ (*left*) and *sadA*$^{-}$ (*right*). (**F**) TIRF images of HSPC300-GFP in *paxB*$^{-}$ (*left*) and *vinA*$^{-}$ (*right*). Substrates were coated with 0 or 30 µg/mL WGA ("Uncoated" or "Low WGA") and 300 µg/mL ("High WGA"). Scale bars, 10 µm.

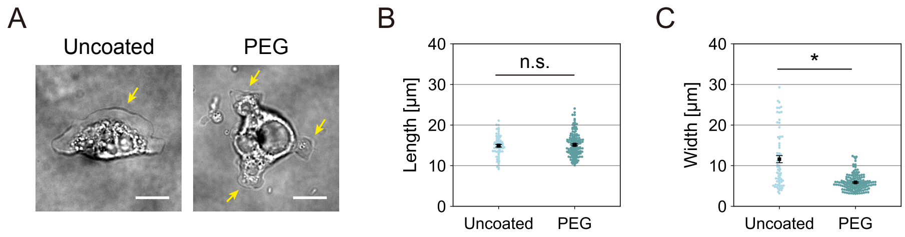


**Figure S8. Cell-substrate adhesion is crucial for protrusion width of *Naegleria gruberi***

(**A**) Transmitted-light images of *Naegleria gruberi* cells on uncoated (*left*) and PEG-coated substrates (*right*). Scale bas, 10 μm. Yellow arrows indicate membrane protrusions. (**B–C**) Length and width of protrusions (mean ± s.e., N = 65 protrusions from 5 cells on uncoated substrates, and 148 protrusions from 6 cells on PEG-coated substrates). Protrusion length was measured as a distance from the cell centroid to the distal end of each protrusion. Student's t-test. **p* < 0.001, n.s., not significant.
